# Deep brain stimulation modulates the dynamics of resting-state networks in patients with Parkinson’s Disease

**DOI:** 10.1101/2020.11.04.368274

**Authors:** Maria T. Gomes, Henrique M. Fernandes, Joana Cabral

**Affiliations:** Department of Physics, NOVA School of Science and Technology, Lisbon, Portugal; Center for Music in the Brain, Department of Clinical Medicine, Aarhus University, Aarhus, Denmark; Life and Health Sciences Research Institute, School of Medicine, University of Minho, Braga, Portugal

**Author notes:** **Correspondence:** Maria Teresa Gomes, Joana Cabral.

**Keywords:** Deep Brain Stimulation, Parkinson’s Disease, fMRI, LEiDA, Dynamic Functional Connectivity, Resting-State Networks

## Abstract

Deep brain stimulation (DBS) of the subthalamic nucleus (STN) is increasingly used for the treatment of Parkinson’s Disease (PD), but despite its success, the neural mechanisms behind this surgical procedure remain partly unclear. As one working hypothesis, it was proposed that DBS works by restoring the balance of the brain’s resting-state networks (RSNs), which is reported to be disrupted in people with PD. Hence, to elucidate the effects that STN-DBS induces on disseminated networks, we analyzed an fMRI dataset of 20 PD patients at rest under DBS ON and OFF conditions. Moving beyond ‘static’ functional connectivity studies, we employ a recently developed fMRI analysis tool, the Leading Eigenvector Dynamic Analysis (LEiDA), to characterize the recurrence of brain-wide phase-locking patterns overlapping with known RSNs. Here, STN-DBS seems to increase the Default Mode Network (DMN) occurrence in individuals with PD. Since the DMN is usually disturbed in PD patients presenting deficits in cognition, our observation might be suggestive that STN-DBS contributes to a normalization of the PD-induced cognitive impairment.

Moreover, we addressed the effects of DBS lead placement on RSNs balance, considering the overlap between the DBS-induced electric field and 3 STN subsections. We found that the Visual Network (VN) probability of occurrence increased proportionally to the electric field-limbic STN overlap. Our finding might be indicative that stimulation of the limbic STN is related to the stabilization of visual symptoms sometimes presented by PD patients, which are usually accompanied by VN disruption.

Overall, this study offers new insights into the fine-grained temporal dynamics of brain states portraying the effects of STN-DBS in patients with PD, while at the same time trying to pave the way to improved planning strategies for this surgical procedure.

## 1 INTRODUCTION

Parkinson’s Disease (PD) is one of the most prevalent neurodegenerative disorders, mainly characterized by motor features, including tremor at rest, rigidity, and postural instability. Besides that, non-motor symptoms such as cognitive impairment, depressive symptoms, sleep disorders, and visual hallucinations are common features of PD, as well [1]. Regarding the treatment of PD, deep brain stimulation (DBS) is now well established, and its success stems from the efficacy in alleviating PD symptoms. Nowadays, DBS appears to be beneficial when used early on to have a more extended treatment window [2], and not only in people for whom oral pharmacotherapy has proven to be insufficient. Moreover, although there is an increasing number of DBS targets, the globus pallidus internus (GPi) and the subthalamic nucleus (STN) are the most popular. Still, the STN is usually the preferred choice for providing a more remarkable improvement in reducing PD’s motor complications [3].

Despite the notable success of STN-DBS, some details in its mechanism of action remain unclear. One hypothesis that is especially attractive to the field of neuroimaging is that DBS may restore the balance of the brain’s resting-state networks (RSNs), which are reported to be disrupted in PD patients [4]. In fact, several functional magnetic resonance imaging (fMRI) studies have shown that people with PD display an altered blood-oxygen-level-dependent (BOLD) functional connectivity (FC) between distinct anatomical regions within large-scale brain networks. Owing to the typical difficulty in executing voluntary movements prevalent in PD, a significant number of studies investigated FC changes within the sensorimotor network. As a result, average FC was shown to be weakened in regions such as the primary motor cortex, premotor cortex, and supplementary motor area in individuals with PD compared to healthy controls [5], [6]. Additionally, Tessitore and colleagues observed that PD patients with freezing of gait had altered FC within the frontoparietal networks sub-serving attentional functions [7]. Interestingly, they reported reduced connectivity in specific areas within the visual network (VN) as well. Meppeling and collaborators detected decreased FC in orbitofrontal and occipital regions of PD patients experiencing visual hallucinations [8]. Furthermore, bearing in mind that the Default Mode Network (DMN) is the most studied RSN in the context of PD, and knowing that this network is intimately associated with cognitive processing, several fMRI studies have searched for FC alterations within this functional network. It has been demonstrated that the connectivity of the DMN is disrupted in people with PD that present cognitive impairment [9]—[11].

Considering the changes in brain FC relating to PD described in the literature, an interest in understanding how STN-DBS affects the PD-disrupted functional networks soon arose. However, until recently, there was no official certificate assuring the safety of acquiring fMRI data in individuals with DBS implants, let alone with the stimulator switched ON inside the scanner [12]. Consequently, only limited work has investigated the effects of STN-DBS in fMRI data in PD patients. In a first study, Jech and colleagues showed that BOLD signals increased in ipsilateral subcortical structures under DBS [13]. Two years later, Stefurak et al. found more distributed signal increases in premotor cortices, ventrolateral thalamus, putamen, and cerebellum in patients undergoing DBS [14]. Besides, Mueller and co-authors used a data-driven and parameter-free analysis method called Eigenvector Centrality mapping and identified increased connectivity in the premotor cortex [15], [16]. Also, Kahan and collaborators employed a dynamic casual modeling approach and showed that DBS was associated with increased coupling of the cortico-striatal, direct, and thalamo-cortical pathways. Moreover, a study conducted upon examining the long-term effects of STN-DBS demonstrated significant local structural connectivity and global FC changes. These changes were observed in regions previously associated with pathological changes in PD, suggesting that DBS helps rebalance the affected networks [17].

After multiple studies showing fMRI acquisitions to be safe and elaborate and modeling animal testing, Medtronic’s Activa^®^ portfolio received in 2015 an extended magnetic resonance conditional CE certificate for full-body MRI [18]. In agreement with this whole-body CE certificate and using a dataset composed of 20 PD patients who underwent STN-DBS, Horn and colleagues characterized changes in the average connectivity of motor brain regions as a function of motor STN-DBS modulation. The results showed that well-placed electrodes led to motor network normalization towards the properties found in healthy individuals. In turn, poorly placed leads did not result in substantial motor network changes [12].

Even so, the great majority of PD- and DBS-related fMRI research conducted so far has focused on FC computed over the whole recording session, implicitly assuming that brain connectivity remains relatively static over time [19], [20]. Despite the great amount of knowledge provided by such ‘static’ FC analysis about the macro-scale organization of brain activity, the characterization of FC over time, or ‘dynamic’ FC (dFC), reveals additional insights to gain a better understanding of brain function. However, the clinical impact of dFC analyses is still being established [21].

Here, we analyze dFC in a cohort of 20 PD patients with STN-DBS [11] using Leading Eigenvector Dynamic Analysis (LEiDA). This method, developed by Cabral and colleagues [22], allows detecting reoccurring states of BOLD phase locking (PL states) that have shown to overlap with known RSNs [20], [23], [24]. This analysis technique allows characterizing phase-locking states in terms of probabilities of occurrence and transition profiles on a subject-by-subject level, thus enabling statistical comparisons between DBS OFF and ON conditions. Additionally, we evaluate the impact of the precise DBS lead location in RSN dynamics by computing the correlation between the weighted electric field-STN overlap and the probabilities of occurrence of the BOLD PL states identified with LEiDA.

## 2 MATERIALS AND METHODS

### 2.1 Study population

#### 2.1.1 STN-DBS cohort

The rs-fMRI dataset used in this work was composed of 20 patients suffering from PD who underwent STN-DBS surgery, and it was previously published [12]. PD patients were 63.7 ± 6.6 years old when they received STN-DBS surgery, and the sample included four females. All patients had the DBS surgery with target STN for idiopathic PD between April 2010 and April 2016, having received two quadripolar DBS electrodes (model 3389; Medtronic). Microelectrode recordings were performed to verify lead placement. The study was carried out in accordance with the Declaration of Helsinki and was approved by the internal review board of Charité – Universitätsmedizin Berlin (master vote #EA2/138/15). More detailed information about the participants is described elsewhere [12].

### 2.2 Magnetic Resonance Image acquisition

#### 2.2.1 Scanning

PD patients were scanned 30 ± 21 months after surgery. Most of the subjects had been implanted at least 11 months at the scan time, except for one subject that was scanned 4 months after receiving STN-DBS surgery. Postoperatively, patients (in medication ON condition, due to logistical reasons) were scanned in a Siemens Magnetum Aera 1.5 T MRI, having received an rs-fMRI scan of approximately 9 minutes under DBS ON condition. Then, patients were briefly taken out of the scanner, for an interval of 5-15 minutes, to turn the impulse generator OFF. Once the PD symptoms reappeared (checked by an experienced neurologist), the rs-fMRI scan was repeated in DBS OFF condition. The scan parameters of MRI images used in this study were: T_1_ MP-RAGE: voxel size 1 × 1 × 1 mm, TR 2200 ms, TE 2.63 ms; rs-fMRI EPI scan: voxel size 3 × 3 × 3 mm, 24 slices with distance factor 30%, A ≫ P phase encoding, TR 2690 ms, TE 40 ms, FOV readout 200 mm. Two hundred and ten volumes were acquired for each condition resulting in a total scan length of rs-fMRI scans of 2 × 9.42 min. Total scan time was slightly less than 30 min, and the B1 + RMS value was always kept below 2.0 μT (conforming to the magnetic resonance-conditional regulations of the Medtronic Activa CE-certificate).

#### 2.2.2 Pre-processing, registration, and parcellation

Rs-fMRI scans under DBS ON and OFF conditions were pre-processed using the Lead-Connectome toolbox (available at http://lead-connectome.org), a complete structural-functional connectomic analysis pipeline in MATLAB (MathWorks, Inc., Natick, MA, USA) which is part of Lead-DBS software (available at http://lead-dbs.org). Briefly, rs-fMRI time series were detrended, and then, motion parameters, mean cerebrospinal fluid, and white matter signals were added as nuisance regressors for noise removal. No global signal regression was performed. The BOLD signals were band-pass filtered between 0.009 and 0.08 Hz. These frequency range limits are traditionally used for preprocessing rs-fMRI data, allowing not only to discard high-frequency artifacts originating from cardiac and respiratory fluctuations but also to focus on the most meaningful frequency range of resting-state signals [22], [25], [26]. Lastly, spatial smoothing with an isotropic 6 mm full width at half-maximum Gaussian kernel was applied. The complete and detailed pipeline is described elsewhere [27]—[29].

The resting-state functional EPI images were coregistered to T1-weighted structural images. Subsequently, the T1-weighted images were coregistered to standard MNI space. Following this and aiming to reduce the dimensionality of the voxelbased data (Voxels × Time), the MNI brain was parcellated and estimated into *N* = 80 different cortical and subcortical noncerebellar brain areas using a custom-made ‘DBS80’ parcellation. The BOLD signals were then averaged over all voxels belonging to each of the 80 brain regions considered, and, with this, data was reduced to size *N* × *T_scan_*. Being *T_scan_* the number of TRs per session, *T_scan_* = 210 for eighteen subjects under DBS ON and OFF, *T_scan_* = 140 for one of the 20 subjects under both conditions, and for another subject, *T_scan_* = 210 TR when DBS was ON and *T_scan_* = 113 TR when DBS was OFF.

### 2.3 Data analysis

#### 2.3.1 BOLD phase dynamics

BOLD phase coherence connectivity [26], [30]—[32] was used to obtain a time-resolved dynamic phaselocking tensor *(**dPL**)* with size *N* × *N* × *T*, where *N* = 80 is the total number of brain areas considered in the current parcellation scheme (‘DBS80’ parcellation) and *T* is the total number of recording frames in each scan, which varies between 111, 138, and 208 (after removing the first and last time points from the total *T_scan_* of 113, 140, and 210, respectively, to exclude the boundary artifacts that the later applied Hilbert transform might induce).

In detail, to compute the phase coherence and obtain a whole-brain pattern of BOLD phase-locking (PL), we first estimated the phase of BOLD signals in all *n* = 1,.*N*brain areas over time *t, θ*(*n, t*), using the Hilbert transform, which expresses a signal *X* in polar coordinates as *X*(*t*) = *A*(*t*) *cos*(0(*t*)), where *A*(*t*) is the instantaneous time-varying amplitude and *θ*(*t*) is the instantaneous time-varying phase. In Figure 1, we have a representation of the BOLD signal phase of one brain area ***n*** over time as ***e^iθ(n,t)^***, where **sin(**θ(*n*, *t*)****) represents the imaginary part of the analytical phase, and cos(θ (*n*,*t*)) represents its real part. More precisely, cos(θ (*n*,*t*)) captures the oscillatory dynamics of the original BOLD signal. Additionally, in Figure 2 A, we represent the portrait of all *N* = 80 BOLD phases during a representative interval of TRs.

**Figure 1.**
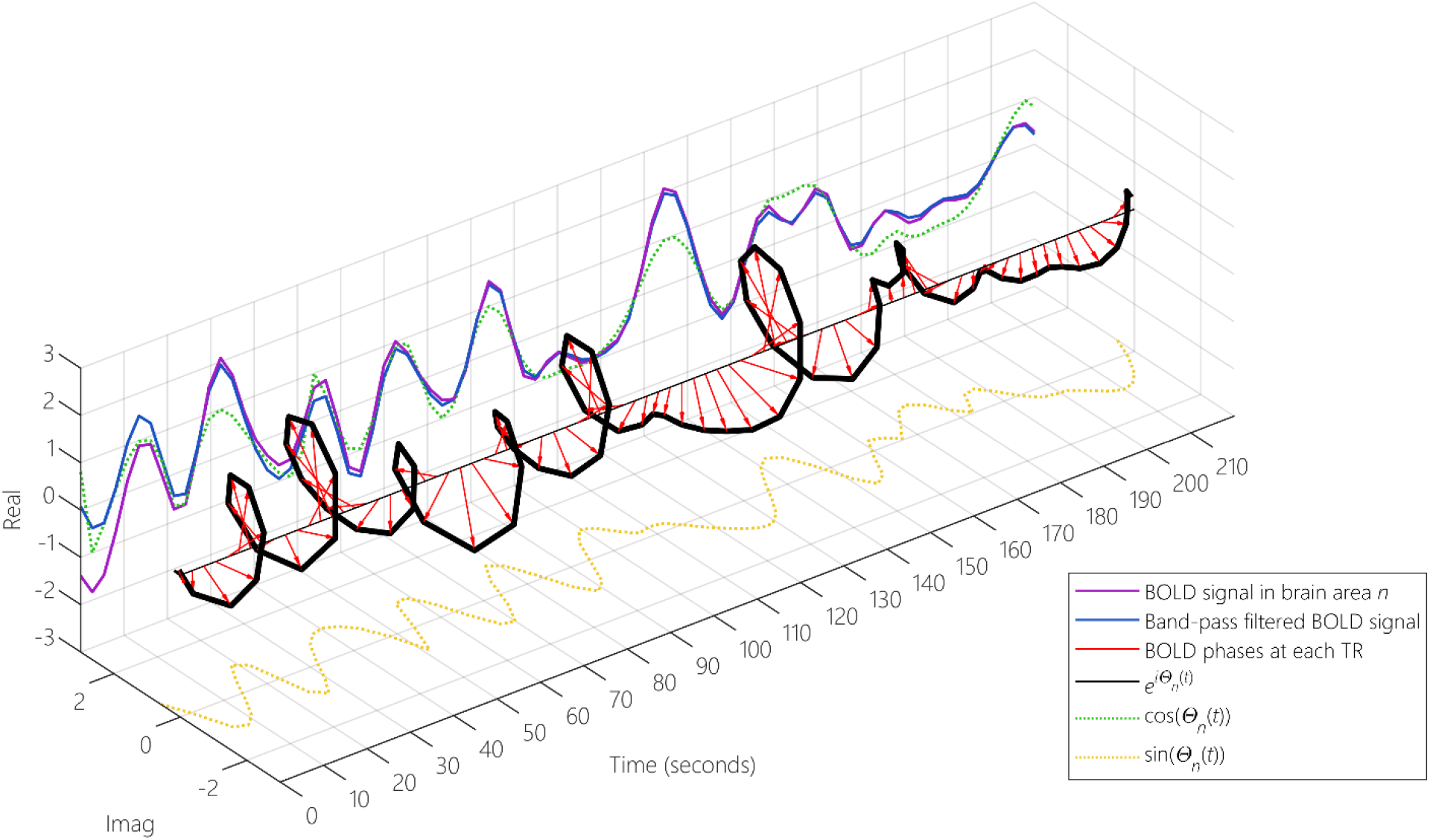
Illustration of the analytical BOLD signal obtained using the Hilbert transform. The BOLD signal of a given area of the brain, ***n***, (purple line), is first band-pass filtered between 0.009 and 0.08 Hz (blue line), and then, using the Hilbert transform, it is transformed into an analytical signal with real and imaginary parts. The phase dynamics of that analytical signal can be represented over time as **e^iθ^_n_** (black line), where the real part (green dashed line) corresponds to **cosθ_n_** and the imaginary part (yellow dashed line) to **sinθ_n_**. The phases of the BOLD signal at each TR are here represented as red arrows.

**Figure 2.**
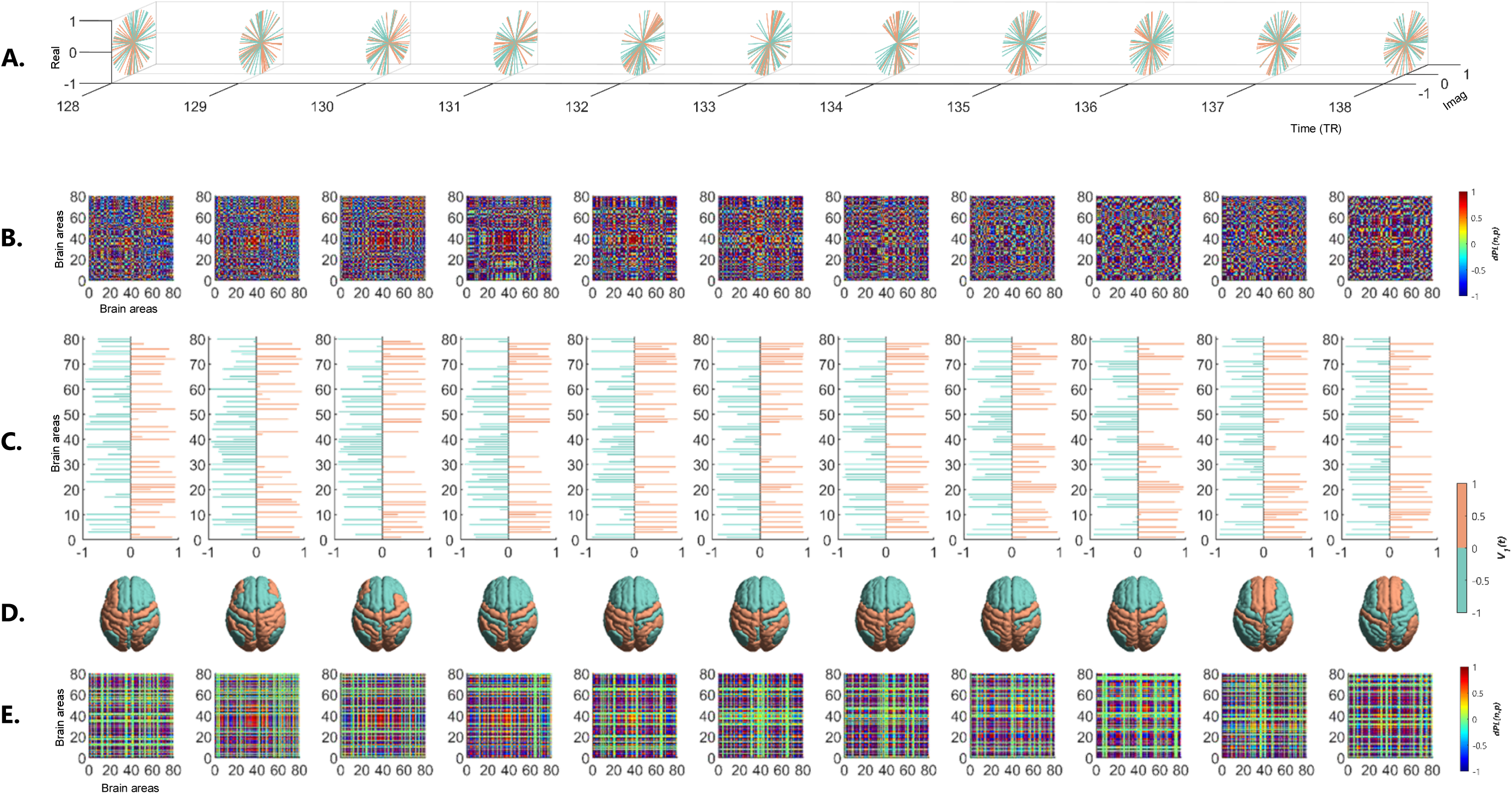
Time-evolving patterns of BOLD phase-coherence obtained with Leading Eigenvector Dynamic Analysis (LEiDA). At each TR (of a representative interval from TR = 128 to TR = 138), (A) the BOLD signal phases in all *N* = 80 brain areas are portrayed as **e^iθ(t)^** where the real and imaginary axes represent the cosine and sine of the Hilbert phase, respectively. (B) The instantaneous phase-coherence between each pair of brain regions can be obtained from the cosine of the phase difference at each time point and are visually represented as phase-coherence matrices, ***dPL(t)***. (C) The instantaneous leading eigenvector, ***V*_1_(t)**, which is the vector that best captures the main orientation of all BOLD phases at that TR (illustrated in (A)), can be obtained from the phase-coherence matrices, ***dPL(t)*** Each element in ***V*_1_(*t*)** corresponds to the projection of the BOLD phase in each brain area into the leading eigenvector and it is colored according to the relative direction with respect to the main one (orange: positive; green: negative). (D) The signs of the elements in ***V*_1_(*t*)** (orange/green) are visualized in cortical space, representing the instantaneous brain phase patterns and illustrating brain areas’ division into two communities (with positive or negative BOLD phase projection into ***V*_1_**(***t***)). (E) The *N × N* dominant connectivity pattern at each TR, captured by ***V*_1_**(***t***), can be retrieved by calculating the matrix product of ***V*_1_**(**t**) with its transpose ***V*_1_**(***t***)^T^.

Given the phases of the BOLD signals, the phase coherence between each pair of brain areas ***n*** and ***p*** at time ***t, dPL(n,p,t)***, was obtained using: ***dPL(n,p,t)*** = **cos(*θ (n,t)* – θ(*p,t*))**. From the result of this equation, three assumptions were made: if two areas ***n*** and ***p*** have temporarily aligned BOLD signals (i.e., with no phase difference) at a given TR, the phase-locking value ***dPL(n,p, t)*** was close to 1 (**cos(0°) = 1**); instead, at time points where the BOLD signals are orthogonal, ***dPL (n, p, t)*** was close to 0 (**cos(90°) = 0**); finally, when the BOLD signals have a maximum phase difference, ***dPL(n,p,t)*** was close to −1 (cos(180°) =—1). With this, it was possible to calculate the ***dPL(t)*** matrix for each time point *t*, where we can visually analyze the coherence in phase between every pair of regions of the brain (Figure 2 B).

#### 2.3.2 Phase-Locking Leading Eigenvectors

Next, in order to identify and then describe recurrent patterns in the (*N × N*) ***dPL(t)***, LEiDA was applied considering, at each time point *t*, only the leading eigenvector, ***V*_1_(*t*)**, that is the one with the largest magnitude eigenvalue [22]. The eigendecomposition of ***dPL(t)*** is calculated at each time *t*, as follows: ***dPL(t) = V(t)D(t)V^-1^(t)***, where columns of ***V(t)*** are the corresponding eigenvectors of ***dPL(t)*** and ***D(t)*** is the diagonal matrix of the eigenvalues of ***dPL(t)***. The first (i.e., largest magnitude) eigenvector, ***V*_1_(*t*)**, is considered to represent the BOLD PL pattern at each time point with size *N* × 1. Notice that since ***dPL(t)*** is symmetric, the respective eigenvectors are orthogonal (***V*^-1^(*t*) = *V*(*t*)^*T*^)** and the eigenvalues are real.

As observed in Figure 2 C and D, the leading eigenvector, ***V*_1_(*t*)**, contains *N* elements (each representing one brain region), and their sign (positive or negative) can be used to separate brain areas into communities according to their BOLD-phase relationship [21], [33]. When all elements of ***V*_1_(*t*)** have the same sign, it means all BOLD phases follow the same direction with respect to ***V*_1_(*t*)**. If instead, ***V*_1_(*t*)** has elements of distinct signs (i.e. positive and negative), the BOLD signals follow different directions with respect to the leading eigenvector, which naturally divides the brain areas into 2 clusters or communities concerning the corresponding sign in ***V*_1_(*t*)**. The magnitude of each element in Vi(t) indicates the ‘strength’ with which the corresponding brain area belongs to the community in which it is placed in. The leading eigenvector ***V*_1_(*t*)** (of dimension *N*× 1) captures the dominant connectivity pattern of ***dPL(t)*** at every time *t*, which can be reconstructed into matrix format *N* × *N* by computing the outer product ***V*_1_(*t*)** ⊗ ***V*_1_(*t*)** (Figure 2 E).

#### 2.3.3 Recurrent BOLD PL states

To explore whether there are specific PL configurations that differentiate PD patients in STN-DBS ON and OFF conditions and achieve a state-based representation, the leading eigenvectors of all 20 patients in both DBS ON and OFF fMRI scanning sessions — corresponding to a total of *T_total_* = 8083 observations with *N* = 80 dimensions each — were clustered into a set of discrete states (or recurrent patterns).

Applying a k-means clustering algorithm to all the 8083 leading eigenvectors, ***V*_1_(*t*)**, across time points, subjects, and conditions, the large number of observations were divided into *k* clusters. Depending on the number of *k* clusters selected, *k* BOLD PL states were returned as solution, with higher *k* revealing rarer, more fine-grained, less frequent, and often less symmetric network configurations. For this study, 19 partitions (with *k* = 2 to *k* = 20 clusters) were considered, having the calculations been repeated 1000 times to ensure the stability in the results. The applied clustering algorithm relied on an iterative process to find the solution that minimized the distance between each (*N* × 1) observation and the closest (*N* × 1) cluster centroid. In this case, we chose to use the Cosine distance for allowing a significantly reduced computation time compared with the Euclidean and City Block distances. According to the number of *k* clusters selected, each observation (or each leading eigenvector) representing a time point was assigned to the cluster (among the *k* clusters in consideration) whose centroid is closest located. As a result, a (1 × *T_total_*) output vector of cluster assignments was obtained, and each cluster can be represented by its (*N*× 1) vector of centroids, ***V_C_***, portraying distinct BOLD PL states.

#### 2.3.4 Between-condition comparisons

To evaluate how the exploration of this repertoire of BOLD PL states returned by LEiDA varied between conditions, we calculated the probability of occurrence (or fractional occupancy) of each PL state, as well as the probabilities of switching from a given PL state to another.

A (non-parametric) permutation-based paired sample t-test (5000 permutations) was used to identify significant differences between DBS ON and OFF conditions, using permutations of group labels to estimate the null distribution, which was then computed independently for each population.

The statistical analyses were performed using MATLAB (version 2019a, MathWorks, Inc., Natick, MA, USA). For statistical significance, it was initially considered a *p-*value of 0.05 or less. To counteract the problem of multiple comparisons, the Bonferroni correction was applied, shifting the *p*-value threshold to *p* < 0.05/*k*.

#### 2.3.5 Correlation between BOLD PL states and reference functional networks

To validate the emergence of well-reported RSNs from the identified BOLD PL states, we here examined if the corresponding centroids shared any spatial organization similarities with seven large-scale distributed brain networks estimated by Yeo and colleagues [34].

First, both the Yeo parcellation mask into seven resting-state functional networks (available at http://surfer.nmr.mgh.harvard.edu/fswiki/Cortical_Parcellation_Yeo2011) and the ‘DBS80’ parcellation (in which our BOLD PL centroids were defined) were taken in the MNI152 space. Then, we calculated, for each of the 80 brain areas (defined in the ‘DBS80’ parcellation), the proportion of voxels assigned to each of the seven reference functional networks. As a result, seven vectors (of dimension 1 × 80) representing the intrinsic functional brain networks in the ‘DBS80’ space were obtained.

Subsequently, all the negative values of the BOLD PL centroids were set to zero so that only areas with positive BOLD phase projection were considered to correlate with the seven reference RSNs. Finally, the correlation between the BOLD PL centroids obtained from our clustering analysis and the seven functional networks was estimated computing the Pearson’s correlation (with associated *p*-values). Importantly, this calculation was done across the entire range of *k* (from 2 to 20).

#### 2.3.6 Impact of the overlap between the DBS-induced electric field and the STN

Given the relevance that the precise location of the DBS lead might have on fMRI brain networks, we considered the overlap between the DBS-induced electric field and three subparts of the STN (motor, associative, and limbic) [12], [35], [36]. These weighted overlap values were then correlated (Pearson’s correlation) with the probabilities of the distinct BOLD PL states identified with LEiDA, searching for significant relationships between electrode placement and specific DBS-induced effects at the brain-network level.

## 3 RESULTS

### 3.1 States of BOLD Phase-Locking affected by STN-DBS

In a first step, we searched for the BOLD PL patterns whose properties most significantly and consistently change when DBS is switched ON. To do so, we compared between ON and OFF conditions, the probabilities of occurrence of all PL states identified with LEiDA across the different partitions into *k* = 2, 3, …, 20. In Figure 3, we report the *p*-values from between-condition comparisons calculated in terms of probability of occurrence for all partitions. Initially, *p*-values were analyzed concerning a standard significance level of 5% (blue dashed line in Figure 3). Subsequently, to take the increased probability of false positives caused by a higher number of comparisons into consideration, the Bonferroni correction was applied, shifting the significance threshold from *p* = 0.05 to *p* = 0.05/*k* (green dashed line in Figure 3). Although 21 solutions passed the standard uncorrected 5% threshold between the probability of occurrence in DBS ON versus OFF conditions (blue circles in Figure 3), only a single PL state (PL state 2 for *k* = 4; green asterisk in Figure 3) survived the correction by the number of clusters within the given partition model (*p* = 0.0076).

**Figure 3.**
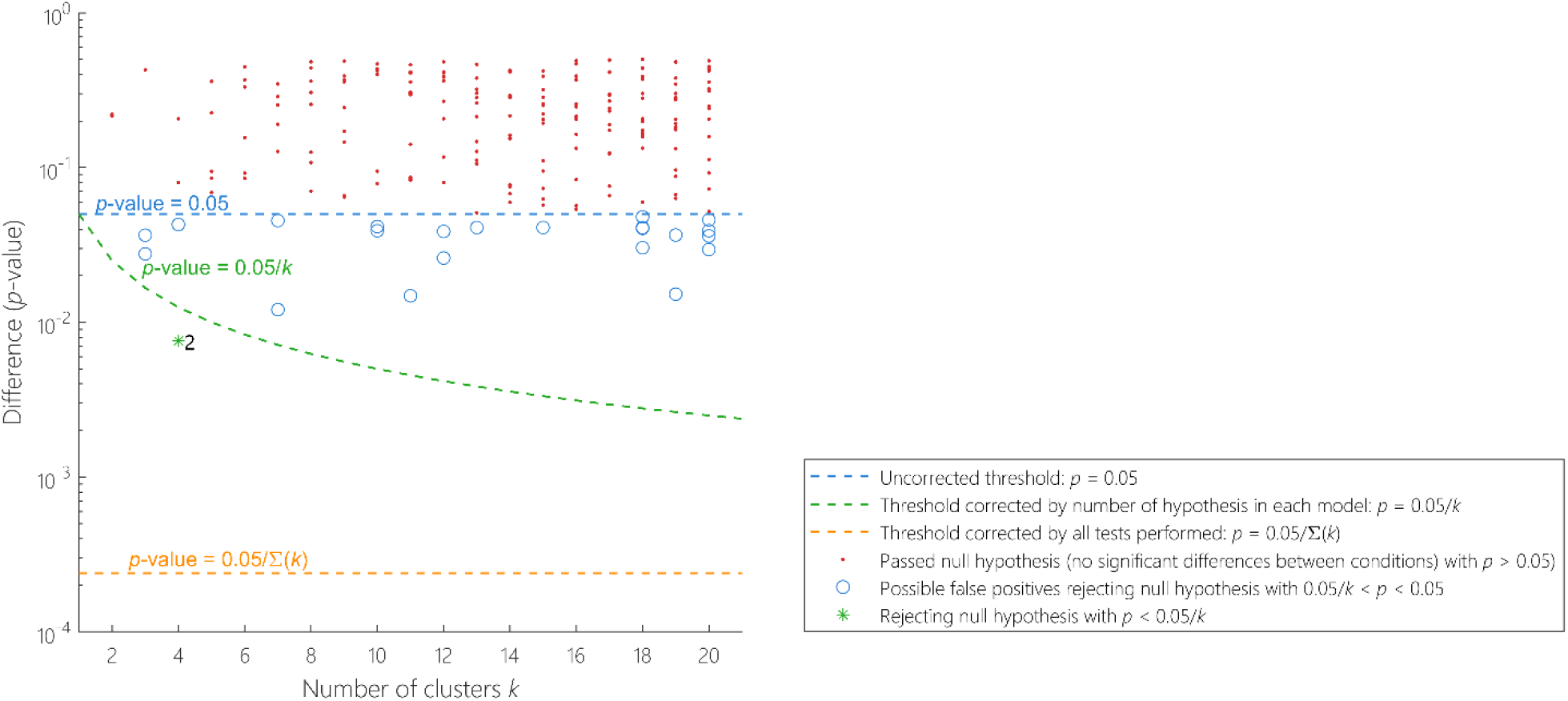
Significance of between-condition differences in phase-locking (PL)-state probability of occurrence as a function of *k*. For each partition of the sample into *k* = 2 to 20 PL states (19 partition models), we plot the *p*-values associated with the comparison between all PL state probabilities of occurrence in DBS ON and DBS OFF conditions. We find that, although most BOLD PL states do not present significant differences between conditions (red dots falling above the *p* = 0.05 threshold, blue dashed line), some PL states show a significant difference in probability of occurrence between DBS ON and OFF conditions (blue circles), passing the standard threshold *(p* < 0.05), but not surviving the correction for the number of clusters in each repertoire of PL states (*p* < 0.05/*k*), henceforth considered false positives. Only one PL state (green asterisk), in terms of probability of occurrence, fall below the corrected threshold by the number of independent hypotheses tested in each partition model *(p* = 0.05/*k*, green dashed line).

We represent in Figure 4 the cluster centroids vectors, *V_C_*, of the PL states with most significant between-condition differences in terms of probability, i.e., with the lowest associated *p*-value, for each of the 19 clustering models considered (with *k* from 2 to 20). Here, areas with the same color (green or orange) have their BOLD phase projected in the same direction as the leading eigenvector.

**Figure 4.**
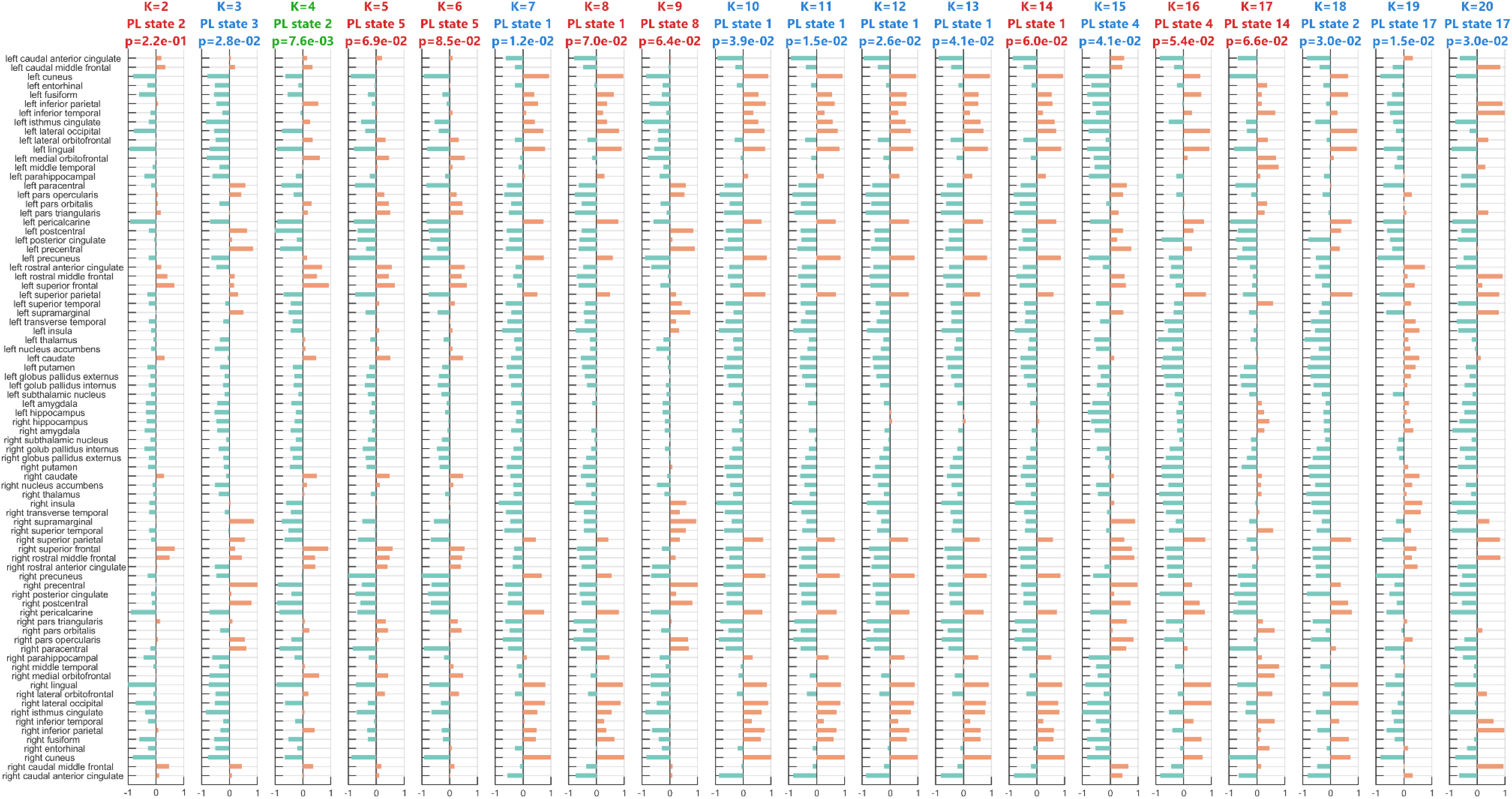
BOLD phase projection, in vector format, of the phase-locking (PL) states with the most significant difference in probability of occurrence between DBS ON and OFF conditions over the range of 19 partition models. After clustering all the 8083 eigenvectors into *k* = 2 to 20 clusters, we identity, for each partition model, the PL state with the most significant difference in probability of occurrence between DBS ON and OFF conditions, that is, with the lowest between-condition *p*-value. Among these 19 PL states, 10 passed the standard threshold of 0.05 (highlighted with a blue title), and only one – PL state 2 for *k* = 4 – survived the Bonferroni correction (highlighted with a green title). These BOLD PL states are here represented by their cluster centroids vector, ***V_C_**, where each element (horizontal bar) represented the BOLD phase, in each of the *N* = 80 brain areas, projected into the corresponding leading eigenvector (captured by the *N* elements in ***V_C_***). The areas *n* whose BOLD phase projection is positive (***V_C_*(*n*)** > 0) are colored in orange, whereas areas *n* with negative BOLD phase projection into the leading eigenvector (***V_C_*(*n*)*** < 0) are colored in green.

### 3.2 Identifying the most relevant BOLD PL state

After analyzing the robustness of the k-means algorithm results and owing to the reliability as a dynamic property that the probability of occurrence has proven to have [21], [23], it was selected for more in-depth analysis the partition into *k* = 4 PL states, as it includes the PL state with the most significant between-condition difference in terms of probability (PL state 2). Moreover, the clustering solution with *k* = 4 ensures a sufficiently detailed subdivision of the BOLD PL configurations and, consequently, reveals meaningful fine-grained structures and networks that differentiate the DBS ON and OFF conditions.

#### 3.2.1 Exploration of the most relevant repertoire of BOLD PL states

The repertoire of PL states returned by LEiDA when choosing *k* = 4 (the number of clusters that allows the best differentiation between conditions in this fMRI experiment) reveals network patterns involving distinct brain subsystems represented in different forms in Figure 5.

**Figure 5.**
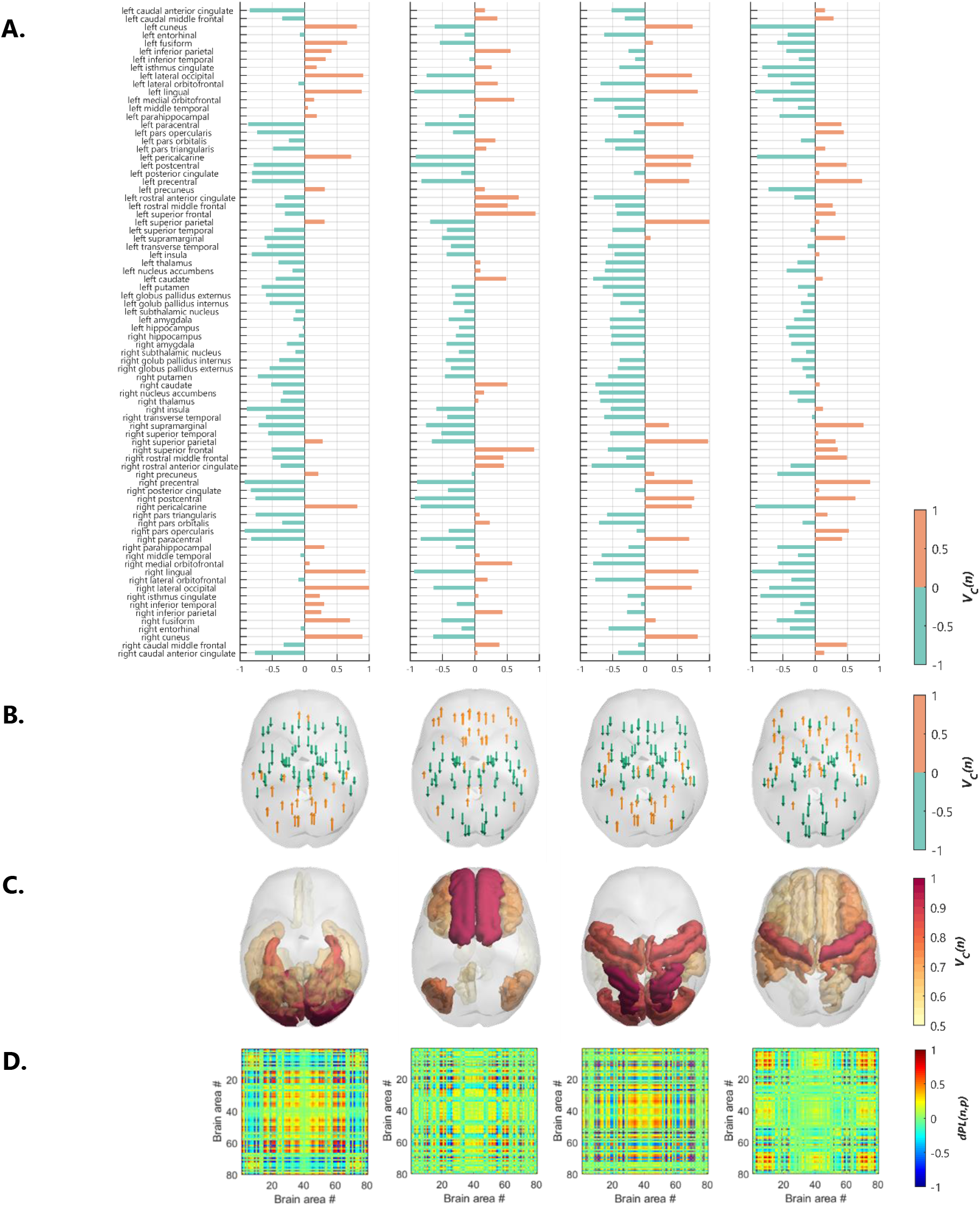
The 4 BOLD PL states more accurately representing the fMRI data from PD patients. A repertoire of 4 recurrent PL states was obtained from clustering the 8083 leading eigenvectors of BOLD phase coherence over time using here *k* = 4. Each PL state is characterized by its **(A)** cluster centroids vector, ***V_C_***, and as **(B)** arrows placed at the center of gravity of each brain area, where the *N* = 80 elements represent the relative projection into ***V_C_*** and are colored in orange when projecting in the main direction of ***V_C_*** and green when projecting in the opposite direction. The brain areas projecting positively into ***V_C_*** are **(C)** rendered in the cortex as patches colored from cream to red according to lower or higher contribution to that brain community, i.e., lower or higher magnitude of (positive) BOLD phase projection. The phase-alignment between each pair of brain regions is captured in the form of an **(D)** 80×80 phase-locking matrix, obtained by computing ***V_C_*** ⊗ ***V_C_***^***T***^.

First, in Figure 5 A, we represent the 4 (*N* × 1) cluster centroids vectors, *V_C_*, representing each BOLD PL state, where each element, ***V_C_(n)***, indicates how the corresponding brain region projects into the leading eigenvector (captured by the *N* elements in ***V_C_***). Thus, orange elements correspond to areas whose projections of the BOLD phase into ***V_C_*** are positive, and green elements correspond to regions with negative BOLD phase projection with respect to the leading eigenvector.

In Figure 5 B, the *N* = 80 vector elements of the 4 cluster centroids, ***V_C_***, are displayed as arrows representing the direction and magnitude of the projection of each brain area into ***V_C_***, which visually divides the brain into two communities according to the referred relative BOLD phase projection. Interestingly, it is easily observed in this representation of arrows in the cortex (as well as in the vector format – Figure 5 A) that these two communities are symmetric across the midline and divide the brain into two distinct brain subsystems.

Furthermore, the 4 PL states are rendered onto a cortical surface (Figure 5 C), coloring only brain areas with positive BOLD phase projection (represented in orange in Figure 5 A and B). The color and transparency of these brain areas are proportional to the magnitude of the corresponding cluster centroids vector’s element. Areas colored in red correspond to higher (positive) magnitude, whereas areas colored in cream correspond to lower magnitude.

Finally, the PL states are represented in the form of 80 × 80 matrices computed as the outer product of the cluster centroids vector (Figure 5 D). Such matrices illustrate the dynamic functional connectivity between each pair of brain areas, where red indicates positive connectivity (in-phase), and blue indicates negative connectivity (out of phase).

For each of these 4 BOLD PL states, we present in Figure 6 the corresponding probabilities of occurrence in both DBS ON and DBS OFF conditions with asterisks representing statistical significance (the values of probabilities (mean ± standard error of the mean) and associated *p*-values are summarized in Table 1). For the selected clustering model (*k* = 4), a significant difference in probability of occurrence between DBS ON and OFF conditions was detected for the PL state 2. This BOLD PL state occurred 22 ± 2 % of the time when DBS was OFF, increasing to 27 ± 2 % when switching DBS ON (*p* = 0.0076). Corroborating and extending on the findings of Figure 3, we identify the BOLD PL state 2 for *k* = 4 as the most relevant PL state, presenting a significant increase in probability when DBS is switched ON in patients with PD.

**Figure 6.**
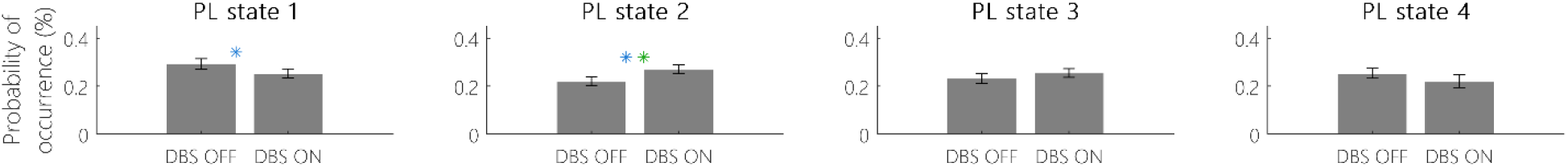
Probabilities of occurrence (mean ± standard error of the mean) of the 4 BOLD PL states most accurately representing the fMRI data from PD patients in DBS ON and OFF conditions. After obtaining the 4 PL states for the clustering solution *k* = 4, we calculated the corresponding probabilities of occurrence. * Significant between-condition difference before correcting for multiple comparisons. ** Significant between-condition difference after correcting for multiple comparisons.

**Table 1.** Probabilities of occurrence of the 4 BOLD PL states most accurately representing the fMRI data from PD patients in DBS ON and OFF conditions.

| PL state | Probability of occurrence (%) |  |  |
| --- | --- | --- | --- |
|  | DBS OFF | DBS ON | *p*-value |
| 1 | 29 ± 2 | 25 ± 2 | 0.0428 |
| 2 | 22 ± 2 | 27 ± 2 | 0.0076 |
| 3 | 23 ± 2 | 26 ± 2 | 0.2067 |
| 4 | 25 ± 2 | 22 ± 3 | 0.0800 |

#### 3.2.2 DBS-affected PL state overlaps with DMN and FPN

Having designated BOLD PL state 2 as the most relevant PL state in our analysis, due to its most significant between-condition difference in probability of occurrence, we henceforth refer to this PL state as ‘DBS-state’.

This dominant BOLD PL state is characterized by a positive BOLD phase coherence between regions including the anterior cingulate cortex, middle and superior frontal gyrus, inferior parietal cortex, isthmus cingulate cortex, orbitofrontal cortex, middle temporal gyrus, pars orbitalis, pars triangularis, precuneus cortex, thalamus, nucleus accumbens, and caudate nucleus (Figure 7 A).

**Figure 7.**
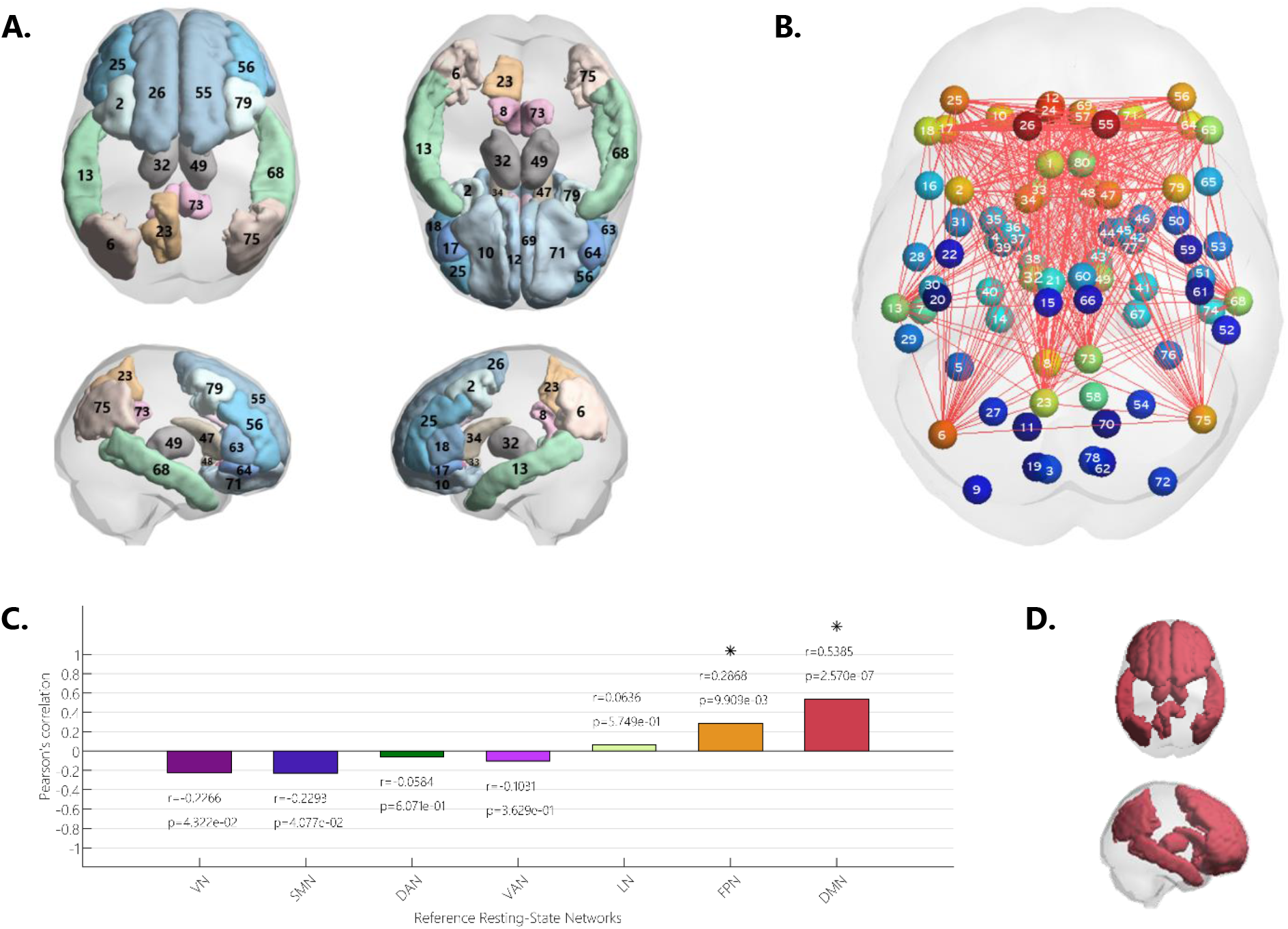
DBS-state: the BOLD PL state with the most significant difference in probability of occurrence between DBS ON and OFF conditions. **(A)** The BOLD PL state 2 (returned by *k* = 4) includes regions with BOLD phase projecting in the same direction as the leading eigenvector, such as the caudal anterior cingulate cortex (1 and 80), caudal middle frontal gyrus (2 and 79), inferior parietal cortex (6 and 75), isthmus cingulate cortex (8 and 73), lateral orbitofrontal cortex (10 and 71), medial orbitofrontal cortex (12 and 69), middle temporal gyrus (13 and 68), pars orbitalis (17 and 64), pars triangularis (18 and 63), left precuneus cortex (23), rostral anterior cingulate cortex (24 and 57), rostral middle frontal gyrus (25 and 56), superior frontal gyrus (26 and 55), thalamus (32 and 49), nucleus acumbens (33 and 48), and caudate nucleus (34 and 47). (**B**) The DBS-state is represented in the cortical space, where functionally connected brain areas (represented as numbered spheres) are linked, and the magnitude values of the BOLD phase projection into cluster centroids vector, ***V_C_***, are used to scale the color of each brain area-sphere. **(C)** This PL state was compared with seven networks of intrinsic functional connectivity by computing the Pearson’s correlation between the corresponding centroids and each one of the seven RSNs used as reference. All Pearson correlation coefficients, *r,* and associated *p*-values, *p*, are reported. * Significant correlation with *p* < 0.05/*k*. VN: Visual Network; SMN: Somatomotor Network; DAN: Dorsal Attention Network; VAN: Ventral Attention Network; LN: Limbic Network; FPN: Frontoparietal Network; DMN: Default Mode Network. **(D)** The DBS-state’s ROIs, which most significantly correlated with the DMN, are rendered onto the cortex and colored in red (color attributed to the DMN according to the color-codes of the reference RSNs estimated by Yeo (see Supplementary Figure S1 B)).

To capture the functional coupling of this PL state’s anatomical areas in the form of a large-scale distributed network, we represent in Figure 7 B the DBS-state in the cortical space, where functionally connected brain areas (displayed as spheres) are linked and colored according to the respective value of the magnitude of BOLD phase projection exhibited in the corresponding cluster centroids vector, ***V_C_(n)*** (see Figure 5 A – PL state 2).

To inspect the functional relevance of this BOLD PL pattern, we overlapped the DBS-state with the seven reference RSNs estimated by Yeo and coworkers [34]. As a result, a significant correlation with the DMN was detected with the highest Pearson’s *r* = 0.5385 and the lowest *p*-value of *p* = 2.5702×10^-7^ (Figure 7 C). Additionally, a meaningful correlation was also observed with the FPN (*r* = 0.2868, *p* = 0.0099). In Figure 7 D, the brain areas with positive elements in the DBS cluster centroid are rendered in the brain, colored in red in accordance with the reference network of intrinsic functional connectivity (from Yeo et al. [34]) that they are mostly correlated to (in this case, the DMN). The seven RSNs used as reference (with corresponding color-code) are illustrated in Supplementary Figure S1 B.

In fact, regions projecting positively into the leading eigenvector in the DBS-state, such as the anterior cingulate cortex [37]-[39], middle frontal gyrus [40], inferior parietal cortex [38], [40], [41], isthmus cingulate cortex [42], orbitofrontal cortex [38], middle temporal gyrus [40], [43], [44], precuneus cortex [37], [39], [40], superior frontal gyrus [45]–[47], and thalamus [37] are reported in the literature as part of the DMN. Moreover, owing to that the DBS-state comprises many frontoparietal areas, including the superior frontal gyrus (coincident with the pre-supplementary motor area) [48], middle frontal gyrus, middle temporal gyrus, inferior parietal cortex [46], [49], and precuneus [46], [47], it makes sense that this PL state also significantly correlates with the FPN.

Furthermore, to portray the variety of RSNs that emerge from the different BOLD PL states obtained with LEiDA, we show in Supplementary Figure S1 A, the *k* cluster centroids (columns) obtained for each partition model into *k* states (rows), rendering and coloring, once again, only the positive elements in the corresponding cluster centroids. Moreover, we color the centroids according to the RSN colorcode from Yeo et al. [34] only when they presented a significant overlap with any of the reference RSNs (surviving a threshold corrected by the number of independent hypotheses tested in each partition model *(p* < 0.05/*k*)). Otherwise, PL states that do not significantly correlate with any reference RSNs were colored in black.

#### 3.2.3 Trajectories between BOLD PL states

Since we are exploring the dynamics of the brain’s activity, we then inspected the trajectories between different PL states by considering the probability of being in a specific PL state, transitioning to any PL state in the following TR. Taking the cluster time courses obtained for *k* = 4, we calculated the probability of switching from a given BOLD PL state to another in each scan and each condition. The average transition matrices, containing the mean probabilities of being in a particular PL state (rows) transitioning to any other PL state (columns), are represented in Figure 8 for both DBS OFF (left) and ON (right) conditions. Notably, the highest probabilities of transition are along the diagonal, representing the probability of remaining in the same BOLD PL state. More specifically, when DBS is ON, the PL state 2 (already identified as the most relevant — DBS-state) is characterized with the highest stability, with a probability of 90% of remaining in that state in the following TR when STN-DBS is switched ON. We have also observed (based on a permutation-based paired t-test) that this self-transition reveals a significant between-condition difference with *p* = 0.0083 < 0.05/*k*. Additionally, a significant increase was detected for the probability of transitioning from the PL state 2 to the PL state 4 when DBS was ON with *p* = 0.0090. Moreover, except for the BOLD PL state 1, which has a minor difference between DBS OFF and ON, all the other 3 PL states present a higher self-transition probability when DBS is ON, compared to when it is OFF.

**Figure 8.**
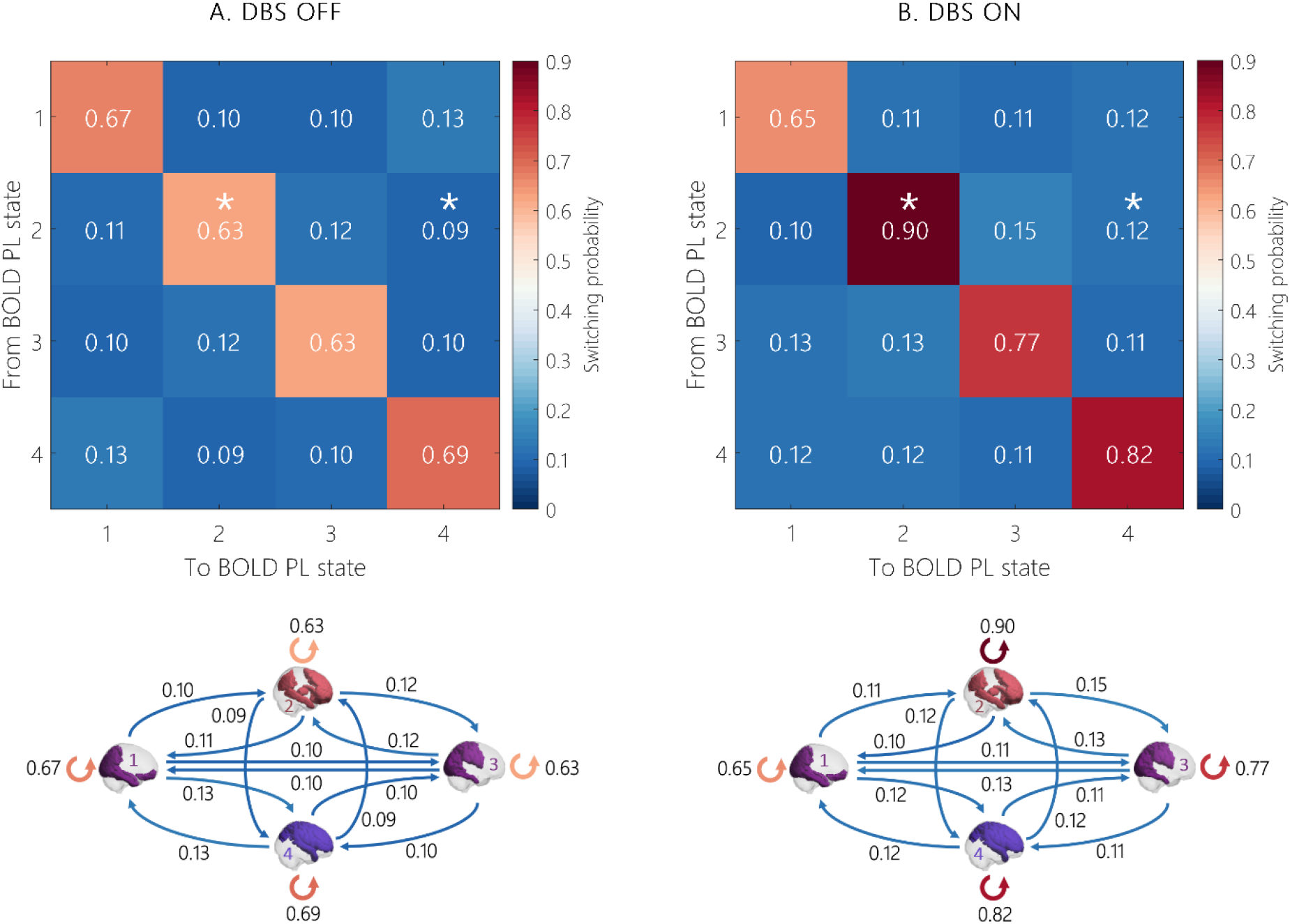
Trajectories of brain activity in state space. For both **(A)** DBS OFF and **(B)** DBS ON conditions, the probabilities of the different trajectories from one of the 4 BOLD PL states to another can be quantitively represented in the form of switching matrices (top) or transition graphs (bottom). * Significant between-condition difference in switching probability.

### 3.3 Effects of electric field-STN overlap on BOLD PL states

#### 3.3.1 Overlap between the electric field and the STN in the DBS-cohort

Based on the estimated overlaps between the DBS-induced electric field and the three-subparts of the STN (motor, associative, and limbic as defined by the DISTAL atlas [156]), we investigated how this weighted overlap modulated the occurrence of the BOLD PL states obtained with LEiDA.

Notice that although these values of overlap between the electric field and the STN correspond to a probabilistic map defining, in some way, the electric field intensity on the STN voxels, they actually reflect better or worst DBS lead placement and consequent STN stimulation.

Since we are here exploring a relationship directly dependent on the *amount* of stimulation in the STN, using the electric field-STN overlap as a regressor to explain phase-locking changes, we have decided to redefine the STN-DBS cohort for this part of the analysis. When analyzing the values of overlap between the electric field and the three STN sub-divisions across the 20 subjects, high heterogeneity was found, and the identification of some subjects with very low overlap between their STN and the DBS-induced electric field led us to establish a minimum threshold of 25% of the mean *overlap* with the total STN.

Accordingly, patients with an overlap between the electric field and the total STN lower than 179 [weighted score] were excluded (see Supplementary Figure S2). Hence, we have elected a final dataset that left out patients 1, 11, and 19, allowing us to preserve, to some degree, the uniqueness of the original fMRI dataset and, at the same time, to exclude those subjects whose electric field-STN overlap was assumed to be insufficient to produce significant changes in whole-brain functional networks.

Furthermore, this redefined dataset led to results very similar to those previously obtained (plot of the correlation-associated *p*-values represented in Supplementary Material as Supplementary Figure S3): the PL state 2 for *k* = 4 was the one presenting the most significant between-condition difference regarding its probability of occurrence, with *p* = 0.0103 that falls below the conservative threshold of *p* = 0.05/*k*. Notably, this result contributed to the validity of the DBS cohort chosen to investigate the effects that the DBS lead placement might induce on BOLD PL states since the exclusion of those three subjects did not return results that go against those obtained in the first part of the study.

This BOLD PL state 2 (for *k* = 4) occurred 20 ± 3 % of the time when DBS was OFF, rising to 24 ± 3 % when switching DBS ON (*p* = 0.0103). Finally, and coincidently to what was detected when LEiDA was applied to the full dataset of 20 patients, this PL state also presented the strongest correlation with the DMN (*r* = 0.5520, *p* = 1.1085×10^-7^), being characterized by a very similar leading eigenvector (see Supplementary Figure S4).

#### 3.3.2 VTA in the limbic STN relates to occupancy of BOLD PL states

Next, we searched for significant relationships between the electric field-STN overlap in each of the three corresponding sub-parts and the probabilities of occurrence, when DBS is ON, of the PL states obtained after applying LEiDA to the dataset excluding patients 1, 11, and 19. To do so, we have calculated the correlation between the DBS-ON probabilities of all the obtained PL states across the 19 partition models and the different values of overlap with the motor, associative, and limbic STN. Plotting the associated *p*-values (Supplementary Figure S5), we found that there was a significant correlation (*p* < 0.05/*k*) exclusively between the electric field-limbic STN overlap and the probabilities under DBS ON of 18 BOLD PL states (the corresponding Pearson correlation coefficients and associated *p*-values are later reported in Figure 9). No statistically significant relationship was detected between the motor or the associative STN overlap, and the PL states’ probabilities of occurrence.

After identifying the BOLD PL states yielding a significant correlation between their probabilities of occurrence and the electric field-limbic STN overlap when DBS was ON, we validated these results by verifying if any significant correlations were detected under DBS OFF (the correlation-associated *p*-values are reported in Supplementary Figure S6). A single PL state was found to have a significant correlation between the DBS-induced electric field and the limbic STN (*r* = 0.5613; *p* = 0.0100) (Supplementary Figure S6 C), meaning that the significant correlations between the weighted overlap and the probabilities of occurrence of the BOLD PL states with DBS ON were not due to artifacts produced by the DBS system.

We further investigated how the overlap between the DBS-induced electric field and the limbic STN modulated the occupancy of BOLD PL states. When DBS was switched ON, we found that the probability of all the 18 PL states exhibiting a significant relationship increased with the overlap values (Figure 9).

**Figure 9.**
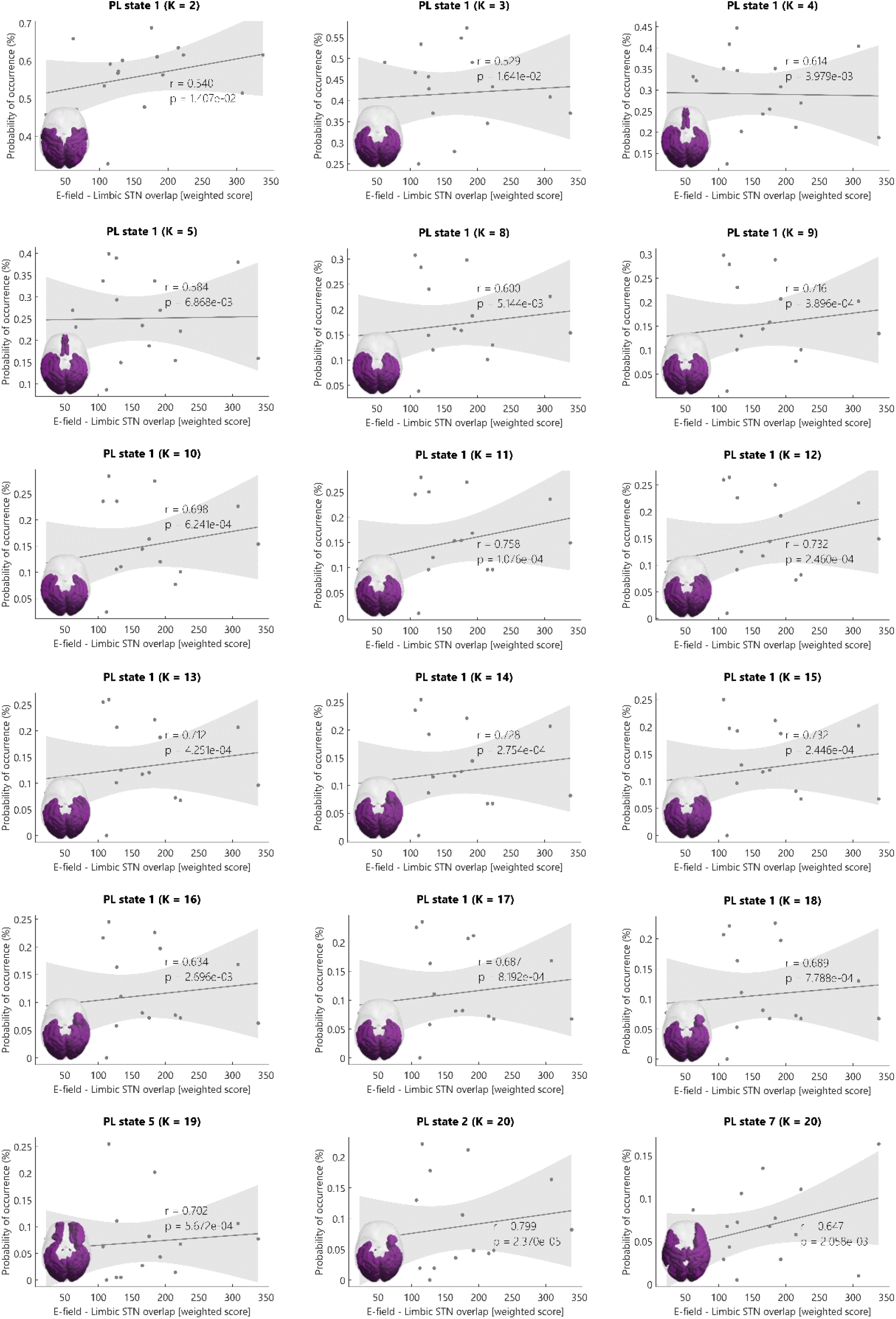
BOLD PL states with significant correlation between the probability of occurrence when DBS in ON and the electric field-limbic STN overlap. 18 BOLD PL states presented a significant correlation between their DBS-ON probability and the values of electric field-limbic STN overlap. For all of them, the probability of occurrence under DBS ON increased proportionally with overlap between the electric field and the limbic STN (Pearson’s *r* and associated *p*-values are reported in the graphs). Moreover, all these PL states correlated most significantly with the VN among the 7 RSNs used as reference (as represented at the left bottom corner of each graph; a more detailed explanation of this result is presented in sub-chapter 3.3.3).

Since the electric field originating from the DBS will overlap with more or fewer limbic STN voxels depending if the electrode is closer or further from the limbic STN, the observed correlation can also be interpreted as the more deviated to the limbic part of the STN the DBS lead is, the higher the occurrence of those 18 BOLD PL states when DBS is ON.

#### 3.3.3 Visual network occupancy correlates with VTA in the limbic STN

Regarding the repertoire of 18 BOLD PL states whose probabilities of occurrence with DBS ON significantly increased in proportion to the overlap between the electric field and the limbic STN, we investigated how they correlate with the reference RSNs from Yeo et al. [34], as described in section 2.3.5. Notably, all the 18 PL states here considered significantly correlated with the Visual Network (VN), as shown in Figure 9, with Pearson’s *r* always above 0.80 and associated *p* < 10^-18^, except for the PL state 7 returned by *k* = 20 (the Pearson correlation coefficients and corresponding *p*-values are reported in Supplementary Table S1).

## 4 DISCUSSION

The present study is one of the first to investigate the effects of STN-DBS on the dynamics of resting-state functional brain networks estimated from fMRI in patients with PD. Representing whole-brain activity at rest as a time-evolving trajectory, we quantify the relative occupancy of a repertoire of whole-brain functional configurations, defined as recurrent BOLD phase-locking patterns over time (PL states), which were validated against pre-established RSNs.

Besides examining the differences of the BOLD PL patterns reoccurring over time during resting-state in patients with PD under DBS ON and OFF conditions, we also quantified the effects of different DBS-lead placements on the occupancy of PL states estimated with LEiDA.

### 4.1 STN-DBS modulates the DMN in patients with PD

Our results revealed a BOLD PL state, the DBS-state, which was much more likely to occur when STN-DBS was ON compared to when it was OFF, meaning that the brain activity in PD patients presented an increased ability to access this PL state under DBS. Moreover, when analyzing the trajectories of brain activity in state space, the self-transitions were most probable; that is, there was a much higher chance for the brain to remain in the exact same BOLD PL state in the following TR (with probabilities varying between 63% and 90%) than switching to a different PL state (with probabilities rounding the 10%). Among these self-transitions, the probability of remaining in the DBS-state was the highest (with a probability of 90%), which reinforces the relevance of this BOLD PL state in the brain dynamics of people with PD undergoing STN-DBS. This result is indicative that the DBS-state is the most stable, such that the brain tends to settle in this PL state when DBS is switched ON.

The DBS-state comprises an extensive network of intrinsic functional connectivity consisting of distinct brain regions commonly reported as part of the DMN such as the anterior cingulate cortex [56], [57], [163], middle frontal gyrus [167], inferior parietal cortex [56], [167], [172], isthmus cingulate cortex [173], orbitofrontal cortex [56], middle temporal gyrus [166]-[168], pars orbitalis and pars triangularis (which form the inferior frontal gyrus) [164], [165], precuneus cortex [57], [163], [167], superior frontal gyrus [169]-[171], and thalamus [57].

#### 4.1.1 Do STN-DBS and consequent DMN modulation restore normal cognition in PD patients?

Owing to the fact that the DMN plays a crucial role in cognitive processing and because the cognitive deficits observed in PD are usually linked to this RSN dysfunction [133], [134], [175], we here propose that the significantly increased probability of the DBS-state when DBS is ON might be indicative of an STN-DBS contribution to the normalization of the cognitive functions that are commonly declined in individuals with PD.

Although PD is mainly associated with degeneration of motor function, studies in patients with this condition suggest that those manifestations are frequently accompanied by cognitive impairment. In fact, while the loss of dopaminergic neurons projecting from the SNpc is particularly severe in the putamen, which explains PD motor symptoms, other dopaminergic projections are affected as well.

Specifically, the caudate nucleus follows the putamen as the second most affected region in dopamine depletion [176]. Moreover, the caudate nucleus is connected to the dorsolateral prefrontal cortex (including the superior and middle frontal gyrus), ventrolateral prefrontal cortex (including the inferior frontal gyrus), and orbitofrontal cortex. Notably, all these last-mentioned brain regions are reported as part of the DMN [177], [178].

Nonetheless, the ventromedial prefrontal cortex also appears to be interconnected with the SN, receiving dopamine innervation, and sending back direct or indirect projections that can modulate dopamine neuron activity [179].

The connection between the caudate nucleus and regions of the frontal cortex is of particular interest because it is widely accepted that the prefrontal cortex plays a critical role in many aspects of cognitive processing [180]. Regarding this, it was hypothesized that cognitive dysfunction in patients with PD would be due to underactivity in the caudate nucleus and, possibly, in the corresponding frontal cortical targets. Remarkably, it was demonstrated that PD patients with executive impairments exhibited significantly less activation of the caudate nucleus, dorsolateral prefrontal cortex, ventrolateral prefrontal cortex, and occipitoparietal cortex, compared with both healthy controls and PD patients with no such deficits [178].

Therefore, as these prefrontal cortical regions receive fibers in a highly ordered topographical fashion from the caudate nucleus, the striatal dopamine depletion effectively interrupts the normal flow of information through such frontostriatal circuits. Hence, the dopaminergic neural death in the caudate nucleus appears to be a serious candidate for mediating the cognitive sequelae of PD [177].

Concurring these assumptions, animal lesion experiments also suggested that the caudate nuclei play an explicit role in cognition, and 18F-dopa PET studies in people with PD have revealed a correlation between dopaminergic depletion of the caudate nucleus and neuropsychological performance, even though these findings have not been universally reported [181], [182].

Interestingly, our results revealed a positive BOLD phase coherence shared between regions such as the superior frontal gyrus, middle frontal gyrus, pars triangularis, pars orbitalis, orbitofrontal cortex, anterior cingulate cortex, and caudate nucleus in the DBS-state. Following the in-phase engagement that these brain areas present when DBS is switched ON, we suggest that the STN-DBS might contribute to the regularization of these frontal areas’ activity, which might lead to the reestablishment of normal cognitive function in PD patients.

From a functional connectivity perspective, PD patients with cognitive impairment commonly exhibit decreased FC within the DMN [183]—[185]. Based on these observations of functional disconnection of the DMN that consequently results in a failure of people with PD to appropriately modulate the DMN activity when facing an executive task, we here propose that the dominance of the DMN under DBS ON is due to a DBS-induced rebalancing of this RSN, potentially leading to improved cognitive performance.

Although the characteristics of DMN dysfunction in PD patients are usually assessed through ‘static’ FC, and we here employed a method based on dynamic FC, interesting parallelisms can be followed. First, it was already observed that areas of the DMN, including the thalamus, anterior cingulate cortex, isthmus cingulate cortex, and superior frontal gyrus exhibited a reduced FC between them in PD patients when compared to healthy controls [186]. On the other side, our results showed that the DBS-state unveiled positive BOLD phase coherence between those DMN-regions (Figure 8.9 — PL state 2 and Figure 8.11). From this observation, we hypothesize that, after STN-DBS, the DMN (which is disrupted by PD) is re-modulated, and the patterns of DMN-connectivity are reestablished in people with PD.

Moreover, a connectivity analysis performed to examine the integrity of the DMN in individuals with PD showing relevant executive deficits reported that, in contrast to healthy individuals, the medial prefrontal cortex and the caudate nucleus were functionally disconnected in PD [134]. As already mentioned, in its turn, the PL state we identified as dominant when DBS was ON is characterized by a positive BOLD phase coherence between the anterior cingulate cortex (which is part of the mPFC) and the caudate nucleus, indicating that these areas are functionally (re)connected. Hence, we suggest that the solid coherence between the BOLD signals derived from those regions observed in the DBS-state might contribute to a normalization of the disrupted (d)FC within the DMN and, consequently, to a potential improvement of the executive deficits underlying the disturbed connectivity between the mPFC and the caudate nucleus caused by PD.

Additionally, the FPN is also thought to be highly relevant for cognition, and its connectivity appears to be reduced in PD patients with mild cognitive impairment [187], [188]. Our results revealed that the extent of the DBS-state also significantly correlated with the FPN. Therefore, the emergence of this RSN in the DBS-state is another argument supporting our hypothesis that the STN-DBS might restore normal cognition in patients with PD.

Regarding the cognitive outcomes across PD patients undergoing STN-DBS, there is quite some heterogeneity in previous studies. Some analyses report deteriorations in performing cognitive tasks when DBS is ON, while others show an outperformance of the PD group with the stimulator switched ON, compared to those who have DBS OFF, and even some exhibit no changes between the two groups. It is crucial to bear in mind, though, that these mixed cognitive outcomes across studies reflect the complexity of the pathophysiological mechanisms of cognitive modifications post-DBS that are individually variable, which contributes to an even more heterogeneity of the results.

Corroborating our assumption that the higher occurrence of the DBS-state (under DBS ON condition) is associated with a restoration of normal cognitive functioning, there are two different studies where an improvement in processing and psychomotor speed with STN stimulation ON compared to OFF was observed [189], [190]. Another analysis that included 48 PD patients reported a significant improvement in psychomotor speed when the stimulator was ON, and an additional subgroup of 15 patients showed a substantial improvement in spatial working memory [191]. In two other studies, a series of 8 and 27 patients, respectively, presented improved memory during ON stimulation condition [192], [193]. Additionally, a study with 18 PD participants described an improvement in one measure of attention with DBS switched ON [194]. Moreover, a report including 7 patients described their better performance in attention, concentration, and frontal executive functions under DBS ON compared to OFF [189]. Likewise, another series of 15 PD patients improved their scores on a test evaluating their executive function when STN-DBS was ON. The authors of the study suggested that those results might be associated with a specific positive effect on the dorsolateral prefrontal cortex (DLPFC) [195] since other reports had already described increases in the activity of the DLPC and cingulate cortex during STN-DBS stimulation [196]. These findings of higher activity in the DLPFC and cingulate cortex support the present study results, where we identified a stronger BOLD phase coherence in regions such as the middle frontal gyrus (area in the human brain where the DLPFC lies) and anterior cingulate cortex.

### 4.2 VN emerges in PD patients with higher stimulation in limbic STN

As mentioned before, there are very different results regarding STN-DBS effects in patients with PD that can be influenced by individual-specific features. In fact, the diversity of neurophysiological outcomes produced by STN-DBS and reported across the existing studies stresses the importance of analyses focused on patient-specific variables, such as the DBS lead placement inside the STN. Concerning different STN-DBS electrodes’ locations, different amounts of voltage are present within the STN, which can be traduced through the overlap between the DBS-induced electric field and the STN.

Building upon a recent study which used the electric field-motor STN overlap as a regressor to explain motor network changes [17], we here investigated the correlation between the electric field-STN overlap within the 3 STN subsections (motor, associative, and limbic as defined by the DISTAL atlas [156]) and the probability of the distinct BOLD PL states’ occurrence when DBS was ON. Exploring this enables us to understand which functional networks are most likely to arise when there is a higher overlap between the DBS-induced electric field and the STN. Our results revealed that the higher the electric field-limbic STN overlap, the higher the probability of activating a BOLD PL state from which emerges the VN.

In PD, retinal dopamine depletion and decreased dopaminergic innervation of the visual cortex can lead to multiple visual deficits, including impairment of visuo-oculomotor control, color vision, contrast discrimination, and visuospatial construction [197], [198]. Besides, individuals with PD exhibiting ophthalmologic problems present alterations in the VN [199]. Specifically, PD patients with visuoperceptual disturbances exhibited decreased average FC in the pericalcarine cortex (where the primary visual cortex is concentrated). Additionally, this region presented reduced FC with the lingual gyrus and cuneus cortex [200]. Nevertheless, decreased FC in visual areas had already been detected in other reports studying connectivity changes in PD [127]. Notably, in our results, the PL states whose probabilities of occurrence increased with the overlap between the electric field and the limbic STN displayed a positive BOLD phase coherence between areas including the pericalcarine cortex, lingual gyrus, and cuneus cortex, among other regions of the visual cortex. This means that these brain areas were functionally connected, in contrast to what was observed in PD patients with visual deficits. Thus, we hypothesize that a higher stimulation of the limbic STN might help functionally reconnect anatomical regions involved in visual processing, potentially improving the visual symptoms of patients with PD.

Remarkably, STN-DBS seems to ameliorate not only motor symptoms but also non-motor symptoms such as visual dysfunction. A study to inspect the effects of STN-DBS on pupillary reactivity and color vision concluded that bilateral STN-DBS significantly improved pupillary constriction size (which is the difference between pupillary area before visual stimulation and its nadir) and color vision as well [201]. Aiming at investigating how STN-DBS alone (i.e., without anti-PD medication) affects oculomotor performance in people with PD, visual features including saccadic and smooth pursuit eye movements were inspected in a different study. Saccadic eye movements are high-velocity, ballistic changes in eye position that bring an object of interest onto the fovea central retinae, whereas smooth pursuit eye movements are tracking movements ensuring that the image of a moving object stays on the fovea. The results showed that STN-DBS significantly improves smooth pursuit and saccade performance in individuals with PD [202]. Similarly, four other studies also reported positive effects of STN-DBS on saccade performance [203]-[206]. Therefore, these distinct researches reinforced the idea that DBS in the STN results in PD patients’ improved ability to perform tasks that rely on visual-motor control.

Additionally, people with PD usually experience turning difficulty, often leading to FOG, and visual information plays an integral role in locomotion and turning sequence [207], [208]. Corroborating the importance of visual processing on FOG, a study has reported that PD patients presenting FOG symptoms exhibit significantly reduced FC within the VN [132]. Furthermore, previous work has already shown that PD patients turn slower and with more steps than healthy controls and that the turn performance is correlated with oculomotor function in a way that individuals who perform later, larger, faster, and fewer saccades turn better [209]. Following this, one study demonstrated that STN-DBS improved turn performance (turn duration), reduced the number of saccades performed during the turns, and increased the amplitude and velocity of the saccade initiating the turn [210]. Thus, it was concluded that STN-DBS contributes to a rebalancing of the PD-induced visual deficits and, as a consequence of that, it also appears to be largely effective in improving turning performance and straight walking hence contributing to the amelioration of FOG symptoms that people with PD may experience.

Our findings of dynamic FC between regions involved in visual processing, which contrast with previous results showing reduced FC within the VN of PD patients with visual problems, combined with the reports of improved oculomotor functions after STN-DBS, lead us to suggest that STN-DBS might help reestablish regular VN connectivity and visual function.

Nevertheless, our observation of increased occurrence of a BOLD PL state from which arises the VN was correlated with a higher overlap between the DBS-induced electric field and the limbic STN. Supporting this, it was already observed in a study where single-unit recordings were obtained from the STN of three monkeys trained to perform a series of visuo-oculomotor tasks that nearly 20% of the recorded neurons became active during eye fixation, saccadic eye movements, or in response to visual stimuli. Besides, it was detected that neurons related to visuo-oculomotor behaviors were primarily found in the anterior ventral-medial tip of the STN, which represents the limbic portion of the STN [211].

Based on all of these, we speculate that placing the DBS lead in a way that the stimulation of the limbic STN is ensured might facilitate VN modulation towards its normal functional architecture, and as a result, there might be an improvement of PD-related visual deficits.

### 4.3 Strengths and limitations

One of our study’s main strengths is the uniqueness of the fMRI dataset, although the lack of healthy controls potentially counts as a limitation. The feasibility of acquiring postoperative rs-fMRI under STN-DBS in a clinical setting has been limited to safety concerns regarding the compatibility between the MRI scan and DBS components. Consequently, very few postoperative fMRI datasets are available. Here, we have a large sample of PD patients and, by functionally neuroimaging them with STN-DBS both ON and OFF, it was possible to non-invasively investigate real-time effects of STN-DBS on the brain’s functional connectivity and fMRI brain networks. Yet, PD patients were scanned in medication ON condition for logistical reasons and to reduce additional motion artifacts, which means that some dopaminergic medication effects cannot be excluded.

Nevertheless, since the field of fMRI under DBS is quite new, this confines the validity of our results, not only because there is little literature that can support our findings but also because the impact of DBS-induced artifacts on rs-fMRI signal has not been thoroughly investigated. In fact, the DBS lead extensions typically produce an artifact in cortical structures. Since these types of fMRI datasets are still new, no software algorithms have been introduced to correct for those artifacts.

Further, using a sophisticated, data-driven analysis technique, LEiDA, we have characterized FC instantaneously and with high temporal resolution. We identified PL states that, even when occurring with relatively low probability, govern the BOLD phase coherence recurrently and consistently across scans and subjects and that were likely to be missed if using analysis over long time windows. This instantaneous nature of LEiDA offers the advantage of detecting more clearly the recurrences of the same PL pattern, thus improving the signal-to-noise ratio in the dFC analysis. Remarkably, in contrast to the most common approaches investigating brain dFC that typically compare dFC patterns by comparing the upper triangular elements of the corresponding ***dFC*** (or ***dPL***) matrices, LEiDA focuses solely on the leading eigenvector of each ***dPL*** matrix, which traduces the dominant connectivity pattern of the multi-dimensional dFC data. This innovative approach not only strongly reduces the dimensionality from *N*(*N*-1)/2 to *N* (where *N* is the number of brain areas considered) while still explaining most of its variance, but it is also more robust to high-frequency noise and thus overcomes a limitation that affects every almost-instantaneous measures of FC [122]. Moreover, identifying recurrent PL patterns allows characterizing BOLD PL states on a subject-by-subject level, whose temporal properties can subsequently be statistically compared between conditions.

Nonetheless, studies of dynamic FC, as it is the present work, also present limitations, mainly because the functional meaning of PL state remains unclear.

We then applied a k-means clustering algorithm to reveal functionally meaningful cluster centroids, which is just one among numerous possible methods that can partition the LEiDA results into significant states. Although this clustering algorithm can be criticized as circular, its implementation is quite simple, and the computational cost relatively low. Succeeding the clustering algorithm-choice issue, there is the selection of the range of clusters. Regarding this, despite having examined partition models exclusively from *k* = 2 to 20 clusters, we demonstrate that results can be robust across a range of partition models, with variances arising from the granularity intrinsic to the number of fixed clusters. Furthermore, the number of clusters selected for subsequent detailed analysis is a trade-off between finer-grained but less robust solutions. Following the approach used in previous work using LEiDA [7], the partition into *k* = 4 clusters was chosen for revealing the BOLD PL state that most significantly distinguished PD patients in DBS ON from DBS OFF conditions, in terms of dynamical properties (in our case, regarding the probability of occurrence). It is worth noting that the goal here was not to identify all the BOLD PL states that occur, but rather to detect those most significantly modulated by STN-DBS; that is, those exhibiting the most significant differences between DBS ON and OFF conditions. However, a partition into a higher number of clusters (and, therefore, states) might be required when addressing certain conditions that affect a specific subsystem optimally defined for higher *k*.

Moreover, concerning the functional relevance of the BOLD PL states identified, a significant overlap was detected between the clusters’ centroids characterizing the different PL patterns and a set of seven resting-state functional networks used as reference [62]. This strong correlation indicates that the BOLD signal phase is sensitive to the same underlying patterns captured using correlation-based analysis techniques, such as ICA. However, LEiDA introduces the advantage of allowing us to observe networks spatially overlapped, which occur transiently at different time frames. Of course, since the methods applied to obtain the PL states and the reference RSNs are different, a perfect match between the centroids and the networks was not expected. Yet, the meaningful overlap indeed validates the functional relevance of the PL patterns, highlighting the strengths of a dynamic analysis of brain-FC.

Another distinctive aspect of our work is the selected parcellation. Unlike all prior studies employing LEiDA, where the Anatomical Automatic Labeling (AAL) atlas parcellation was used, we here elected the ‘DBS80’ parcellation. Although this parcellation has the asset of having been designed specifically for DBS studies, it also lacks validation, which possibly limits our study. The use of this parcellation was also influenced by the fact that for being a coarser parcellation than AAL, it might reduce the theoretical bias induced by the DBS lead extension artifact. Finer parcellations would probably run into the risk of having single parcels totally filled with the artifact.

Notably, this work has also addressed an aspect repeatedly neglected in fMRI DBS context: the impact that distinct DBS electrode placement and hence, different overlaps between the DBS-induced electric field and the STN have on brainconnectivity profiles. Incorporating the electric field-STN overlap into the dynamical analysis of the influence that STN-DBS has on distributed brain networks makes one of the key strengths of the current study since this has rarely been explored and might provide novel insights into the unclear mechanisms of action of DBS.

However, these *overlap* values were calculated using a weighted over a binary model, i.e., the overlap between the electric field and the STN is calculated through the weighted sum over the electric field distribution within the STN domain. In the past, the few studies investigating the effects that the precise DBS lead placement induces at the brain-network level used the volume of tissue activated (VTA) as a regressor. Nevertheless, this VTA has been modeled as a binary region around the electrode, which means that it represents the total amount of tissue stimulated, disregarding if there are areas more activated than others. On the contrary, using the weighted overlap between the DBS-induced electric field and the STN, the degree of tissue modulation is taken into account [17]. Still, additional research is needed to corroborate the feasibility of using this weighted overlap as a predictor of effects on fMRI brain networks.

One final limitation that applies for careful interpretation of our results is that the PD patients included in the current study were not submitted to specific tests evaluating cognitive or visuo-oculomotor functions, which are those here proposed to be positively affected after STN-DBS. The suggestions and conclusions made upon our findings must be cautiously construed, bearing in mind that these are mere speculations based on literature analysis and require further validation to draw inferences in the clinical domain.

## 5 CONCLUSION

Overall, employing the novel LEiDA tool to investigate instantaneous dynamic FC, this study provides a new framework for detecting resting-state functional network specificities associated with STN-DBS. Our finding of increased ability presented by PD patients with STN-DBS to access and remain in a specific BOLD PL state from which emerges the DMN might be indicative that STN-DBS acts by restoring DMN dysfunction. Thus, as this RSN normally reappears as a baseline functional network at rest, it might potentially contribute to a normalization of commonly PD-induced deficits in cognition. Moreover, considering different DBS lead placement as a critical factor for the effects produced by STN-DBS in fMRI brain networks, we detected a higher probability of VN emergence when there was a larger overlap between the DBS-induced electric field and the limbic STN. Again, this possibly means that STN-DBS facilitates the amelioration of non-motor symptoms, in this case, oculo-visual dysfunction. In conclusion, although there is evidence that STN-DBS reshapes restingstate brain connectivity, not many research pieces have been done in that matter, and this work stresses the importance of exploring the underlying post-DBS changes at the brain-network level. Doing so offers insights for further understanding of the potential neuro-modulatory mechanism underlying DBS of the STN and its therapeutic effects in PD, which might contribute to better planned DBS surgeries for targeting the improvement of patientspecific symptoms.

