## Supplementary Material for "Deep brain stimulation modulates the dynamics of resting-state networks in patients with Parkinson’s Disease"

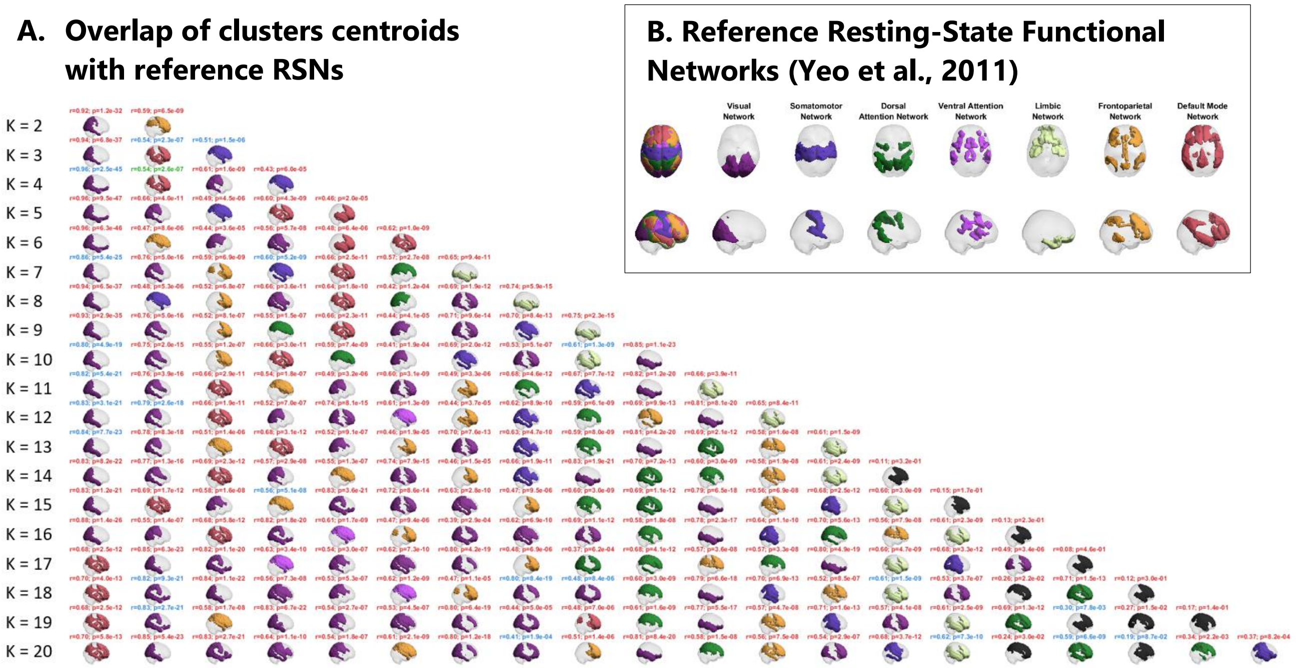


Figure S1 - Overlap between the centroids of the PL states and the reference resting-state functional networks. (A) Representation of the centroids obtained for each k-means clustering solution with *k* ranging from 2 to 20. The centroids of each PL state are represented in cortical space, having selected as ROIs only those with the BOLD phase projecting positively with respect to the leading direction. In turn, the ROIs are colored according to the reference RSN (shown in Supplementary Figure S1 B) with which they most significantly correlate. In case there is no significant overlap with any of the brain networks used as reference, the corresponding centroids are colored in black. Pearson's *r* and associated *p*-value are reported as a title in red when the *p*-value related to the difference of probability of occurrence in DBS ON vs OFF condition of the represented BOLD PL state is greater than 0.05 ($p > 0.05$), in blue when falling below a standard significance threshold of 5 % ($0.05/k < p < 0.05$), and in green when the *p*-value survives a conservative threshold of $p < 0.05/k$, to correct the number of clusters in each partition model. (B) Seven resting-state networks estimated by Yeo using correlation-based FC on 1000 healthy subjects at rest were used as reference to evaluate the correlation with the obtained PL states and are illustrated in top view and side view.

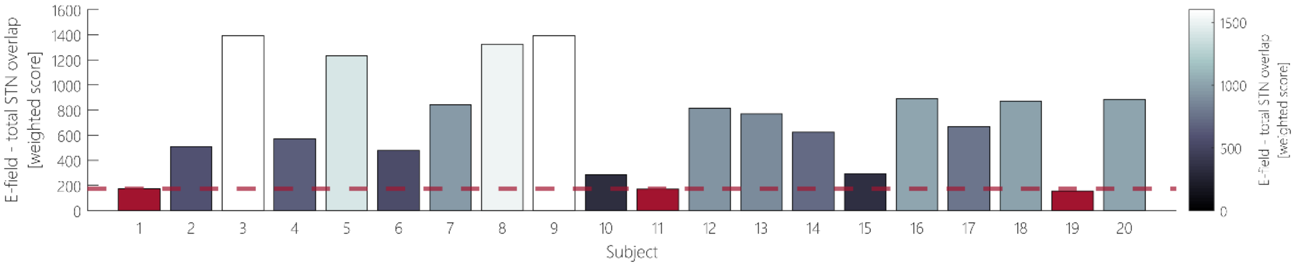


**Figure S2 - Overlap between the electric field and the STN.** Across all the 20 PD patients, it was estimated the overlap between the electric field and the total STN. The colors from black to white indicate a smaller or higher weighted overlap. After defining a minimum threshold of 25% of the mean electric field-STN overlap (here represented as a dark red dashed line), subjects 1, 11, and 19 (with overlap values represented as red bars) were excluded for the analysis of the effects that different DBS lead placements produce on the BOLD PL states identified with LEiDA.

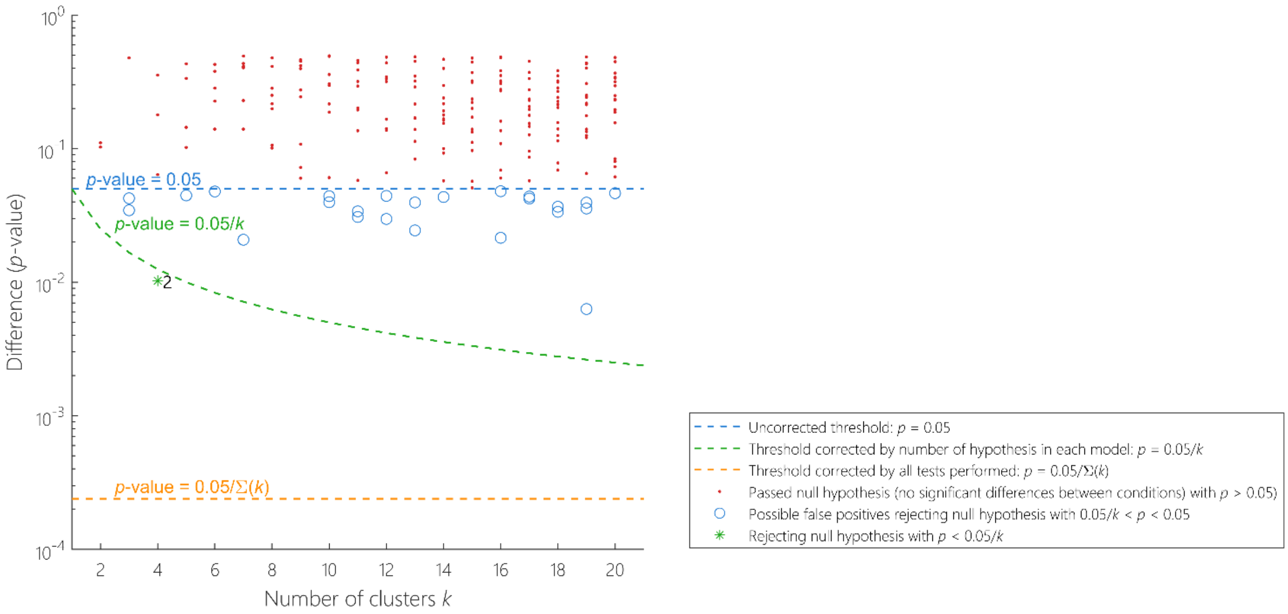


**Figure S3 - Significance of between-condition differences in phase-locking (PL)-state probability as a function of *k* (for a dataset without subjects 1, 11, and 19).** For each partition of the sample into $k$ = 2 to 20 PL states (19 partition models), we plot the $p$-values associated with the comparison between all PL state probabilities of occurrence in DBS ON and DBS OFF conditions. We find that, for this dataset excluding subjects 1, 11, and 19, the only BOLD PL surviving the corrected threshold by the number of clusters ($p < 0.05/k$; green dashed line) and exhibiting a very significant between-condition difference in probability was the PL state 2 returned by $k = 4$ (see Figure 3).

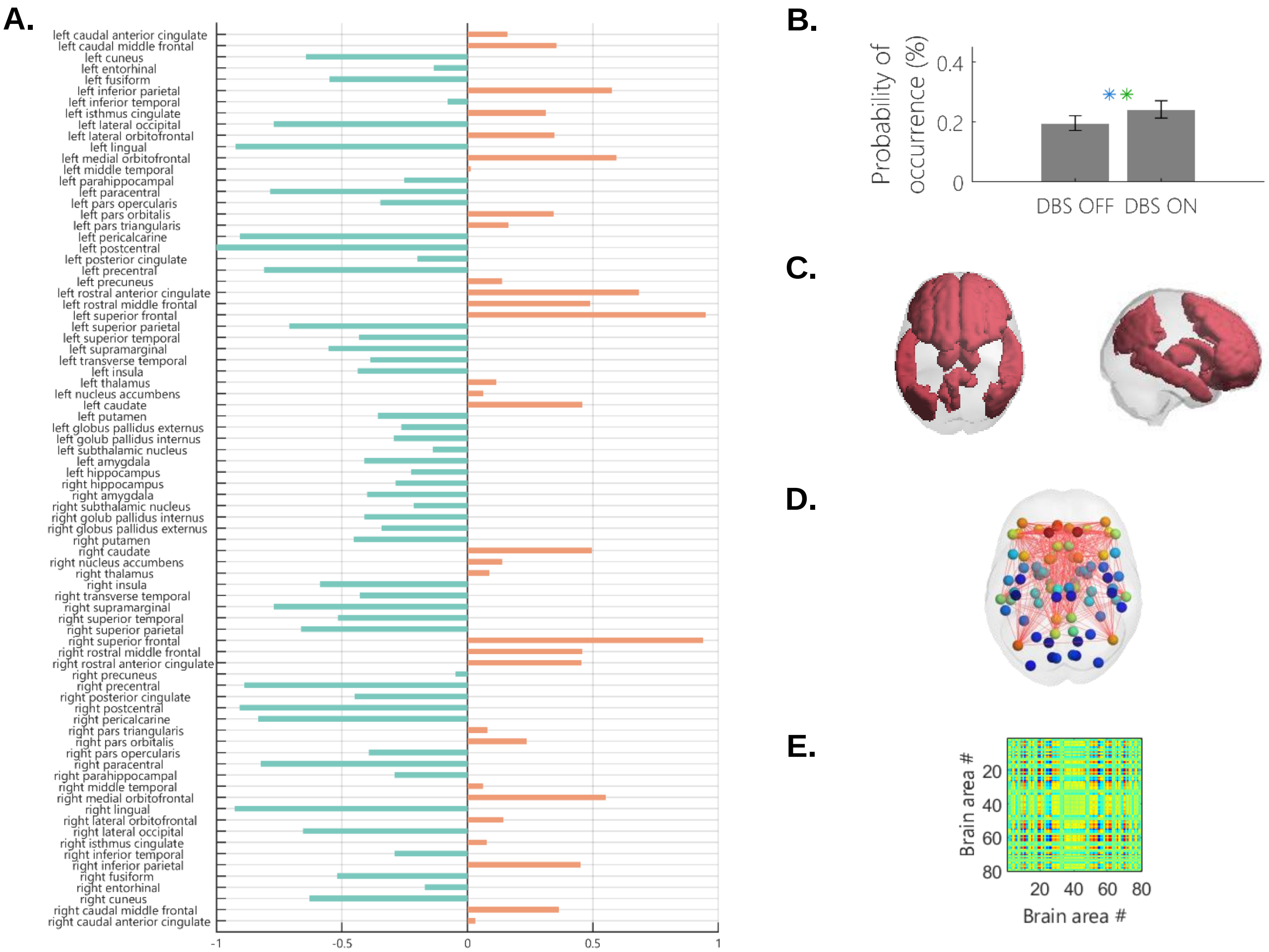


**Figure S4 - Phase-locking (PL) state with the most significant between-condition difference in probability of occurrence (for dataset without subjects 1, 11, and 19). (A)** The contribution of 80 different brain areas to the dominant PL state is represented through its cluster centroids vector, $\boldsymbol{V_C}$. Bars in orange represent areas with positive BOLD phase projection and bars in green represent the brain's regions with negative projection into the leading eigenvector (captured by the *N* elements in $\boldsymbol{V_C}$). **(B)** Differences in probability of occurrence of this BOLD PL state between DBS OFF and ON conditions (20 ± 3 % vs. 24 ± 3 %, respectively, $p$ = 0.0103). ** Significant between-condition difference after correcting for multiple comparisons. **(C)** The brain areas of the most significant BOLD PL state, which most significantly correlated with the DMN, are rendered on the cortex and colored in red (color attributed to the DMN according to the color-codes of the reference RSNs estimated by Yeo (see Supplementary Figure S1 B). **(D)** The dominant BOLD PL state is represented in the cortical space, where anatomical regions (displayed as spheres) that are functionally connected are linked through red lines. The spheres are colored according to the BOLD phase projection's magnitude of the corresponding brain area. **(E)** This significant PL state is also represented as an 80 × 80 matrix computed as the outer product of the centroids vector, $\boldsymbol{V_C}$, which embodies the BOLD phase coherence of each pair of brain areas. The synchronization colored from blue, passing to green, and ending in red corresponds to a negative, null, and positive synchronization, respectively.

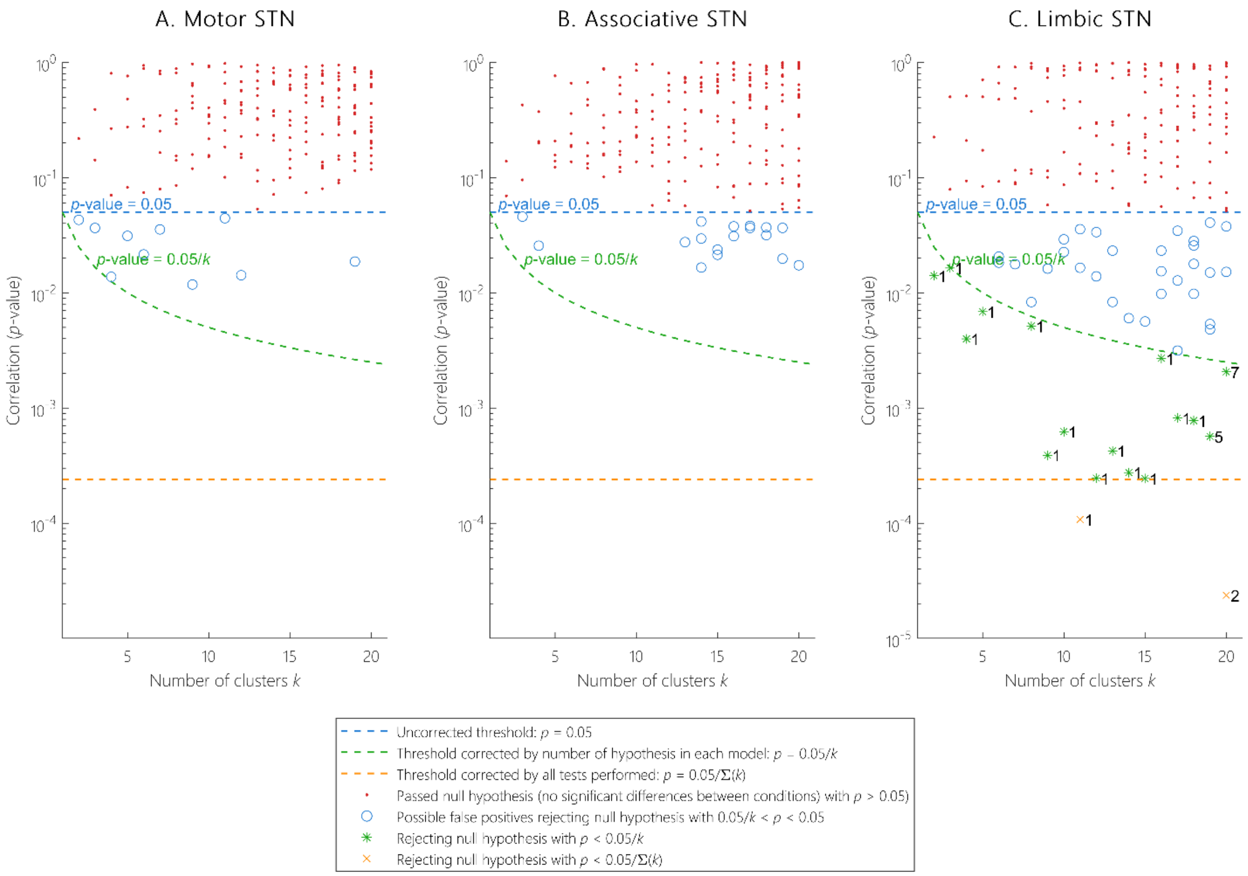


**Figure S5 - Significance of correlation between the electric field-STN overlaps and the probabilities of occurrence under DBS ON condition of the distinct BOLD PL states.** For each partition of the sample into $k$ = 2 to 20 PL states (19 partition models), we plot the $p$-values associated with the correlation between all PL states' probabilities when DBS was switched ON and the overlap between the DBS-induced electric field and the **(A)** motor, **(B)** associative, and **(C)** limbic STN. We observe that for the correlation between the electric field-STN overlap in both motor and associative STN and the probabilities of occurrence of all the PL states, the associated $p$-values either pass the null hypothesis (red dots above the $p$ = 0.05 threshold, blue dashed line) or pass the standard threshold of $p$ = 0.05 (blue circles). However, they never survive the correction for the number of independent hypotheses tested in each partition model ($p$ < 0.05/$k$; green dashed line). On the other side, a great number of BOLD PL states reveal a significant correlation of their probabilities of occurrence with the different values of overlap between the electric field and the limbic STN, with $p$-values falling below the corrected threshold (green stars). Even two PL states surviving the threshold corrected by all tests performed (yellow dashed line) with $p$-values under 0.05/$\boldsymbol{\Sigma}$($k$) (yellow crosses).

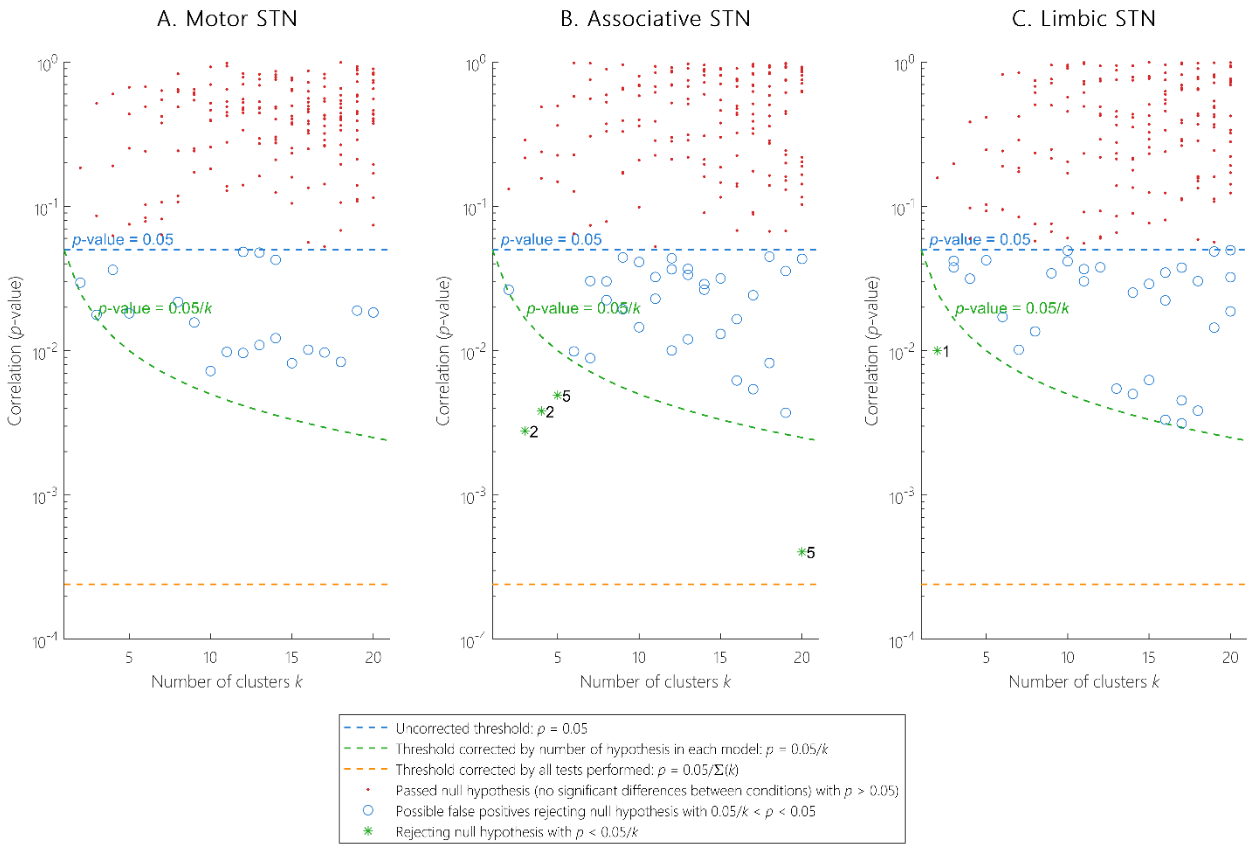


**Figure S6 - Significance of correlation between the electric field-STN overlaps and the probabilities of occurrence under DBS OFF condition of the distinct BOLD PL states.** For each partition of the sample into $k$ = 2 to 20 PL states (19 partition models), we plot the $p$-values associated with the correlation between all PL states' probabilities when DBS was OFF and the overlap between the DBS-induced electric field and the **(A)** motor, **(B)** associative, and **(C)** limbic STN. We observe that for the correlation between the the electric field-STN overlap in the three parts of the STN and the probabilities of occurrence of all the PL states, most of them either fall above the standard threshold of $p$ = 0.05 (red dots) or fall bellow this (blue circles). This observation reveals no significant correlation between the probabilities of occurrence and the electric field-STN overlaps (only 5 PL states – green asterisks – survive the correction by the number of clusters within the given partition model).

Table S1 – Statistical analysis of the relationship between the VN and BOLD PL whose DBS-ON probabilities of occurrence are modulated by the electric field-limbic STN overlap.

| Number of clusters | PL state | Pearson's $r$ | $p$-value |
|---|---|---|---|
| $k = 2$ | 1 | 0.9115 | $8.0283\times10^{-32}$ |
| $k = 3$ | 1 | 0.9366 | $2.8787\times10^{-37}$ |
| $k = 4$ | 1 | 0.9674 | $2.6426\times10^{-48}$ |
| $k = 5$ | 1 | 0.9700 | $1.1967\times10^{-49}$ |
| $k = 8$ | 1 | 0.9461 | $6.0680\times10^{-40}$ |
| $k = 9$ | 1 | 0.9051 | $1.0683\times10^{-30}$ |
| $k = 10$ | 1 | 0.8207 | $1.1817\times10^{-20}$ |
| $k = 11$ | 1 | 0.8540 | $7.7686\times10^{-24}$ |
| $k = 12$ | 1 | 0.8566 | $4.0612\times10^{-24}$ |
| $k = 13$ | 1 | 0.8384 | $2.9297\times10^{-22}$ |
| $k = 14$ | 1 | 0.8071 | $1.5540\times10^{-19}$ |
| $k = 15$ | 1 | 0.8493 | $2.4206\times10^{-23}$ |
| $k = 16$ | 1 | 0.8775 | $1.3201\times10^{-26}$ |
| $k = 17$ | 1 | 0.8870 | $6.7766\times10^{-28}$ |
| $k = 18$ | 1 | 0.8784 | $9.8752\times10^{-27}$ |
| $k = 19$ | 5 | 0.8287 | $2.3519\times10^{-21}$ |
| $k = 20$ | 2 | 0.8904 | $2.1931\times10^{-28}$ |
| $k = 20$ | 7 | 0.5465 | $1.5755\times10^{-7}$ |
